# The Structure of the Lujo Virus Spike Complex

**DOI:** 10.1101/2024.03.13.584793

**Authors:** Maayan Eilon-Ashkenazy, Hadas Cohen-Dvashi, Sarah Borni, Ron Shaked, Rivka Calinsky, Yaakov Levy, Ron Diskin

## Abstract

Lujo virus is a human pathogen that emerged as the etiology agent of a deadly viral disease in Africa. While it is a member of the *Arenaviridae*, it is a distinct virus that does not classify with the classical ‘Old World’ or ‘New World’ groups of viruses in this family. It further utilizes neuropilin-2 (NRP2) as an entry receptor, a property that is not shared by other arenaviruses. So far, structural information is limited to the receptor binding domain of LUJV, and the overall organization of the trimeric complex, as well as the way NRP2 is recognized in the context of the complete viral spike, were unknown. Here, we present the cryo-EM structure of the complete, native, membrane-embedded spike complex of LUJV. We found that NRP2 is bound at the apex of the spike in a way that allows each trimer to engage with a single NRP2. Also, the complete receptor binding site is quaternary, depending on interactions contributed by neighboring protomers. Recognition of NRP2 involves an overlooked arginine-methionine interaction, which we have now characterized. This LUJV’s spike structure, which is the second determined structure of a complete arenaviral spike, points to similarities and differences in the structures of these viral spikes, informing vaccine design and allowing us to be better prepared to combat future outbreaks of this virus.

## Introduction

Zoonotic viruses that circulate in animal hosts and can be transmitted to humans pose a substantial risk to human health, raising a great concern for their potential to cause outbreaks. One such virus is the Lujo virus (LUJV), a contagious and highly fatal pathogen that emerged in a limited but deadly outbreak in South Africa ^1^. No specific treatment is currently available against LUJV, and there is an urgent need to develop effective countermeasures. LUJV was found to belong to the *Arenaviridae* ^1^, which is a large family of enveloped, bi-segmented, ambisense RNA viruses. The mammarenavirus genus, which includes all arenaviruses that infect mammals, is geographically and genetically classified into the Old World (OW) and the New World (NW) groups that are mostly endemic to Africa and the Americas, respectively. Both of these groups contain viruses that can infect humans, like the OW Lassa (LASV) and lymphocytic choriomeningitis (LCMV) viruses, as well as the NW Junín, Guanarito, Sabiá, and Machupo (MACV) viruses. Infection by these viruses may result in mild disease, but it could also develop into severe, often fatal, hemorrhagic fever disease. Interestingly, LUJV is genetically distinct from both the OW and NW groups of mammarenaviruses ^1^. LUJV also differs in the cell surface receptor that it utilizes for cell entry; To attach to and enter their target cells, the OW and some NW viruses use a special glycan modification, termed matriglycan, that specifically modifies α-dystroglycan (α-DG), and most NW viruses use transferrin receptor 1 (TfR1) ^2,3^. The LUJV, on the other hand, exploits neuropilin-2 (NRP2) to attach and enter its target cell ^4^, and it is the only mammarenavirus so far known to do so.

Regardless of the cell entry receptor usage, all mammarenaviruses have trimeric class-I glycoprotein spike complexes that mediate the attachment of the viruses to their cells and ultimately also fuse the viral and host cell membranes ^5,6^. Unique to Arenaviruses, these spikes function as three-partite complexes; the spikes are translated as glycoprotein precursor proteins (GPCs), which are processed by a signal peptidase (SPase) and a subtilisin kexin isozyme-1/site-1 (SKI-1) protease ^7^, yielding a unique structured signal peptide (SSP), glycoprotein-1 (GP1), and glycoprotein-2 (GP2). The SSP, which is unique to Arenaviruses, is unusually long (*i.e.*, 58 aa), could be added in *trans,* and is critical for the spike’s trafficking as well as for its ability to get proteolytically mature by SKI-1 cleavage and fuse membranes ^8–14^. GP1 serves as the receptor-binding domain, and GP2, which is a transmembrane domain, serves as the fusogenic module of the spike ^6,15–17^.

So far, structural information for receptor recognition by an intact, functional arenaviral spike complex is available only for LASV ^17^. Structural information is also available for the recognition of TfR1 and NRP2 by isolated GP1 domains of the MACV and LUJV spike complexes, respectively ^15,16^. However, the structure of the LASV spike complex revealed an intricate, quaternary-dependent organization of the SKI-1 recognition motif that ultimately forms the receptor binding site for matriglycan ^17^. Quaternary-dependent binding of NRP2 was also suggested in the case of LUJV ^16^, but in the absence of structural data for the LUJV spike complex, the mere existence of such quaternary-enabled interactions or their nature is unknown. In fact, it is not known how the trimeric organization of spike complexes differs among distinct mammarenaviruses.

Here, we present the cryo-EM structure of a full-length, membrane-embedded spike complex of the LUJV. This structure reveals the organization of the spike in the membrane as well as the architecture of its ectodomain. Using our previously determined crystal structure of NRP2/GP1 complex ^16^, we leverage our findings to reveal how the NRP2 receptor is recognized in the context of the complete spike complex. Being the second complete arenaviral spike whose structure is available so far, it provides an opportunity to reveal structural elements that are common to arenaviruses as well as unique attributes that are virus-specific. This information will be important for designing future vaccines.

## Results

### The structure of the LUJV spike complex

To decipher the structure of the LUJV spike complex, we expressed the entire spike in mammalian cells with a Flag-tag fused to its C-terminus. We solubilized the cells’ membranes in detergents, purified the spike, and determined a three-fold symmetric, 2.8 Å resolution structure using single-particle cryo-EM (Extended Data Fig. 1, Extended Data Table 1). All three domains that result from proteolytic cleavage of GPC (Fig. 1a) are visible in the density map and are part of the final model (Fig. 1b), which consists of the spike’s trans-membrane and ectodomain but not the cytoplasmic portion. The overall architecture of the spike resembles that of the Lassa virus spike complex ^17^. In the membrane, the spike forms a six-helical bundle made of three central helices of GP2 surrounded by three helices of SSP (Fig. 1b). The central three GP2 helices are wrapped around each other such that each helix penetrates the membrane underneath a neighboring GP2 protomer (Extended Data Fig. 2a). This is achieved by a long helix (α4, Fig. 1b & Extended Data Fig. 2a) that extends to a neighboring GP2 and closely interacts at a kink that forms between the neighbor’s α4 and α5 GP2 helices (Extended Data Fig. 2b). The interaction is facilitated by sandwiching Glu386 and Tyr381 from each of the α4 helices between Thr374/Lys378 and Gln385/Ile390 on the neighboring α4 helices, respectively (Extended Data Figs. b & c). Above the surface of the membrane, the GP2-GP2 interaction is mainly driven by hydrophobic interactions that helix α3 makes with a groove between helix α2 and a preceding loop of an adjacent GP2, as well as electrostatic interactions between two oppositely-charged patches (Extended Data Fig. 3).

**Figure 1.**
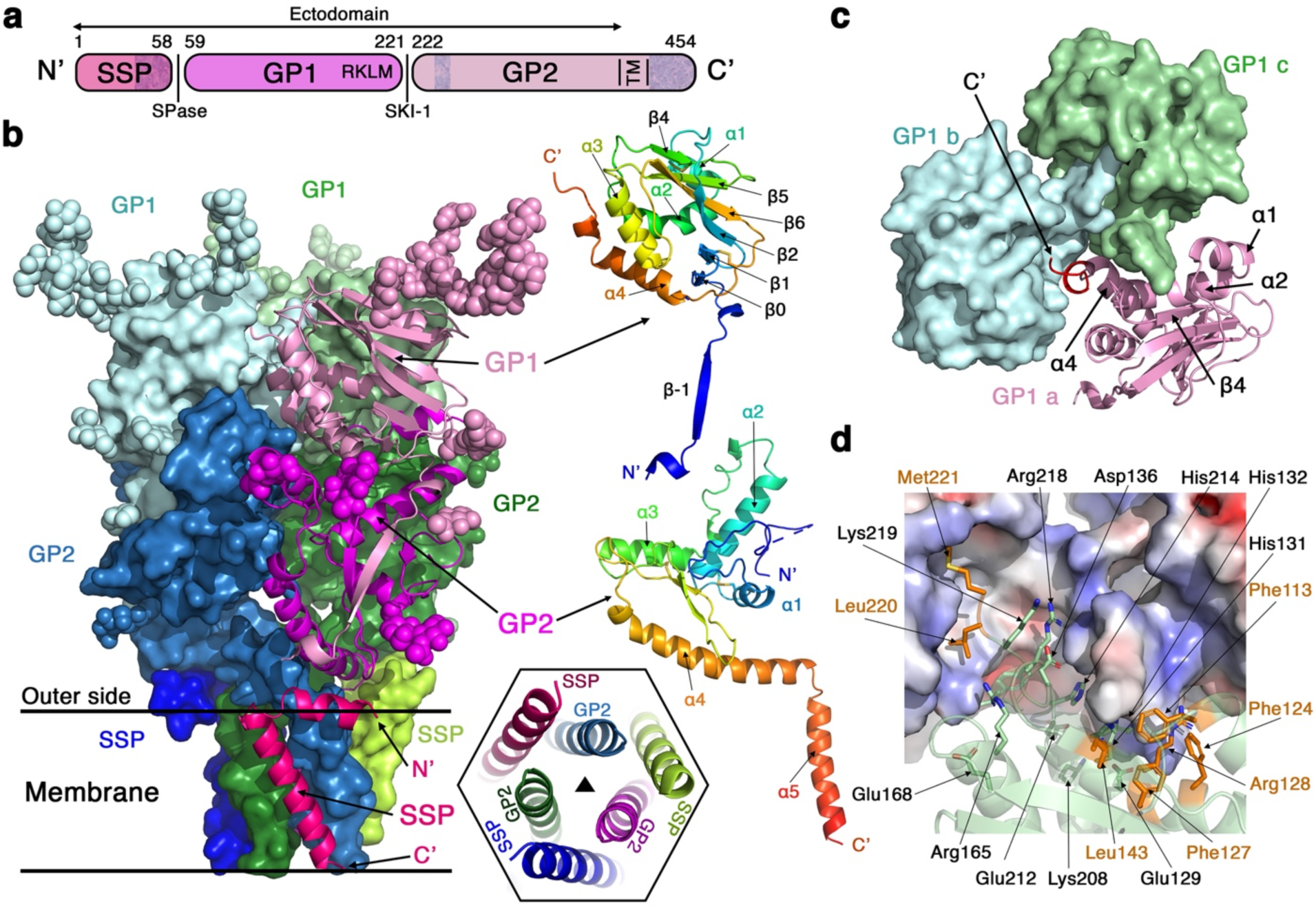
Overall structure of the LUJV spike complex. **a**. Schematic diagram showing the domain partitioning of the spike. Shaded regions indicate parts that are missing from the final model. **b**. The LUJV spike complex is presented using a ‘side’ view. The three GPC protomers are colored pink, blue, and green; for each protomer, the SSP, GP1, and GP2 domains are indicated using different tones. One protomer (pink) is shown in a ribbon diagram, and the organization of its secondary structure is indicated for the GP1 and GP2 domains in two ribbon representations, which are rainbow-colored from their N- to the C-termini. The naming of secondary structure elements in GP1 preserves and extends the naming in PDB ID: 6GH8. N-linked glycans are shown as spheres. The estimated location of the membrane is indicated. The inset shows the organization of the transmembrane helices, presented along the three-fold symmetry axis of the spike (indicated with a triangle) and shown from the ‘bottom’ of the spike. **c**. The interaction between the GP1 domains. A top view shows only the GP1 domains, using the same coloring scheme as in ‘b’. The main GP1 elements that mediate the interaction are indicated. **d**. A close-up view of the interface between the GP1 domains. Two GP1 domains are shown using a surface representation colored by their electrostatic potential (red -5 kT/e; blue 5 kT/e). Key interacting residues are sown as sticks and indicated with arrows. Residues that mediate hydrophobic interactions are highlighted in orange.

The SSP starts with a short helical segment that is parallel to, and partially embedded in, the membrane, followed by a trans-membrane helix (residues 15 to 34) (Fig. 1b). This configuration indicates a topological rearrangement of the C-terminal part of SSP from the outer/luminal side to the inner/cytoplasmic side following cleavage by the signal peptidase (SPase) (Extended Data Fig. 4). The previously observed ^17^ structural role of SSP in stabilizing the spike complex seems to be conserved as the SSP interacts in the membrane with two adjacent GP2 helices, such that it prevents them from unwinding. The SSP itself is held fixed with respect to the trimer by interactions that its short helical N’-segment makes with GP2 (Fig. 1b).

The apex of the spike is stabilized by mutual interactions between the GP1 domains (Fig. 1c). The C-terminus of GP1, resulting from cleavage by the cellular protease SKI-1, together with the preceding helix α4, forms a latch that extends toward and interacts with the neighboring GP1 near the three-fold symmetry axis of the spike (Fig. 1c). This latch-mediated interactions create deep groves between the GP1 domains (Fig. 1c). The tip of this latch is made from the two hydrophobic residues of the ‘RKLM’-SKI-1 recognition site (Fig. 1a & 1d). These hydrophobic residues fit into a hydrophobic pocket on the neighboring GP1 made by Phe124, Phe127, Leu143, and the aliphatic portion of Arg128 (Fig. 1d). The rest of the interacting surfaces are very polar and include Arg218 and Lys219 from the ‘RKLM’ motif, as well as several histidine residues like His131, His132, and His214 (Fig. 1d). Glu212 is also at the interface (Fig. 1d). This glutamic acid points into the symmetry axis of the trimer where it chelates a metal ion (Extended Data Fig. 5a). Although Glu212 likely stabilizes the spike, chelating a metal ion in this site is not required for the spike’s proper function, as mutating Glu212 (*i.e.*, E212A or E212Q) results in spikes that are as efficient as the WT spike in promoting cell entry (Extended Data Fig. 5b).

### Recognition of NRP2 by the LUJV spike complex

We previously determined a crystal structure of an isolated, partially truncated LUJV GP1 domain in complex with its NRP2 receptor ^16^. Superimposing this isolated GP1 domain on the GP1 domains from the trimeric spike reveals almost identical structures with root-mean-square deviation (RMSD) = 1.2 Å, without any apparent conformation changes (Fig. 2a). This observation paves the way to use the information from the GP1/NRP2 crystal structure to understand how the trimeric spike recognizes NRP2. The most noticeable aspect of this interaction is that NRP2 is bound at the apex of the spike (Fig. 2a) with an orientation that overlaps the spike’s three-fold symmetry axis (Fig. 2b). This observation reveals that each spike complex can interact only with a single NRP2 molecule at a time.

**Figure 2.**
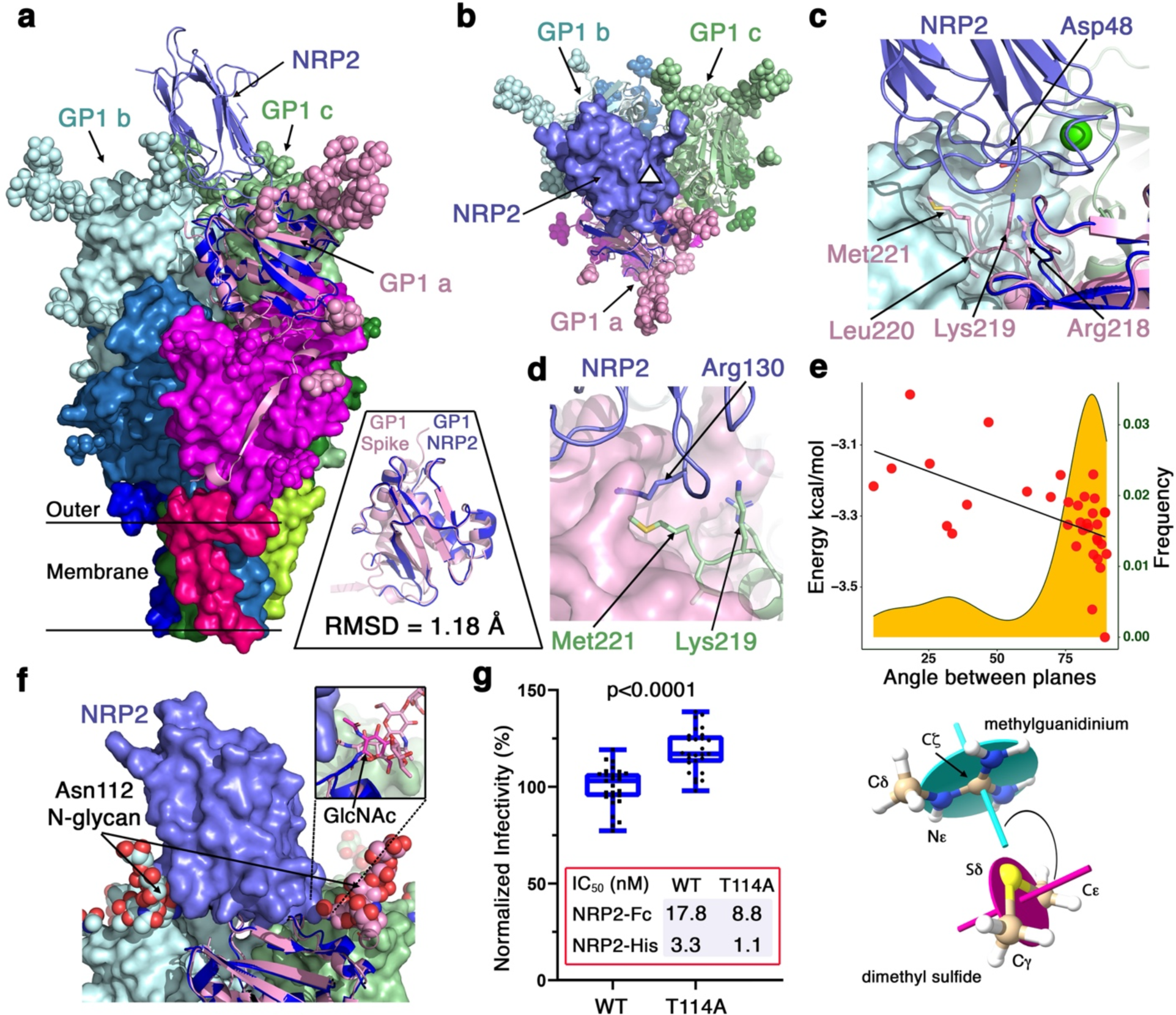
Recognition of NRP2 by the LUJV spike complex. **a**. NRP2 binds at the apex of the spike. The inset shows the superimposition of two GP1 domains from the structure of the spike (pink) and the structure of the NRP2/GP1 complex (blue, PDB ID: 6GH8). The RMSD value based on a common set of 110 Cα atoms is indicated. The spike complex is shown in a ‘side’ view using surface representations for all chains except a single GP1 domain (pink) that uses a ribbon diagram. The coloring scheme is the same as in Figure 1. The NRP2/GP1–spike superimposition places the NRP2 (purple ribbon diagram) at the apex. **b**. A ‘top’ view of the spike. Bound NRP2 overlaps with the spike’s three-fold symmetry axis, which a triangle indicates. **c**. Lys219 from the same GP1 domain that NRP2 binds is positioned such that it can form a salt bridge with Asp48 of NRP2. **d**. Arg130 of NRP2 serves as a lid that closes on the side-chain of Met221 from a nearby GP1. **e**. Methionine’s sulfur atom/guanidinium interaction. The orange chart shows the preferred angle between the guanidinium and the Cψ-S8-Cε planes, as defined in the lower image, for arginine/methionine interactions based on statistical data derived from the PDB (n=663). The red dots show calculated energy values (kcal/mol) for the bond strength between methylguanidinium and dimethyl sulfide as a function of the angle between the planes. A linear fit for the data points (R^2^=0.33) is shown. **f**. NRP2 fits between two N-linked glycans on Asn112 residues from two separate GP1 domains. Inset shows a magnified view of the glycan from the cognate GP1 domain that binds the NRP2. In the NRP2/GP1 crystal structure, the first N-Acetylglucosamine is tilted with respect to the same sugar in the apo-spike structure. **g**. Normalized infectivity of pseudotyped viruses bearing either the WT or a T114A mutant of the LUJV spike complex. Dots represent technical replicates (n=27). The p-value (two-tailed Student’s t-test) for the difference is indicated. Whiskers indicate the minimum and maximum values. Central lines indicate means, and boxes show the interquartile ranges. Results for a representative experiment out of two independent repeats are shown. Below the graph, IC_50_ values for neutralization by NRP2-Fc, and NRP2-His are indicated.

On top of the various Ca^2+^-dependent interactions that a single LUJV GP1 domain forms with NRP2 ^16^, superimposing the GP1/NRP2 crystal structure on the spike’s structure suggests additional quaternary structure-enabled interactions. Namely, Asp48 of NRP2 is perfectly positioned to form electrostatic interaction with Lys219 of its cognate GP1 (Fig. 2c). This Lys219 is pre-positioned by the interaction that the hydrophobic-latch makes with a neighboring GP1 domain (Figs. 1c & 2c). Also, another Lys219 from a neighboring GP1 can form an additional interaction with the main-chain carbonyl of NRP2’s Arg130 by adopting a different rotamer (Extended Data Fig. 6).

A particularly interesting quaternary-enabled interaction involves Arg130 of NRP2. This arginine residue makes important polar interactions with GP1 ^16^. In the context of the trimeric spike, the side-chain of Arg130 further forms a lid that covers the hydrophobic cavity to which Met221 from a neighboring GP1 is inserted (Fig. 2d). In this configuration, the aliphatic tail of Arg130 makes Van der Waals interactions with the side-chain of Met221. Interestingly, this configuration also positions the sulfur atom of Met221 (Sδ) below the plain of Arg130’s guanidino group (Fig. 2d). The contribution of a sulfur-guanidino interaction to the stability of the complex is not clear as this kind of interaction was not described before. Bioinformatic analysis of unique 34,376 structures at a resolution better than 2.2 Å from the PDB reveals n = 771 instances of methionine’s sulfur atoms that are at interaction distance from guanidino groups of arginine residues (3.2 Å – 3.8 Å from the Cζ atom of arginine, which is the closest atom) (Fig. 2e). This set was split into interactions at polar or hydrophobic environments (*i.e.*, nearest water molecule is at a distance smaller, or greater than 3.5 Å from the Cζ atom of arginine and the Sδ atom of methionine). For n = 663 interactions that are in a less polar environment, there is a preference for orientations that minimize the Sδ - guanidino group distance while maximizing the distances of the methionine’s Cψ and Cχ atoms (*i.e.*, the angle between the normal of the guanidino group’s plane and the normal of the Cψ-S8-Cχ plane approach 90°) (Fig. 2e). To determine the interaction energy, we employed Double-Hybrid Density Functional Theory (DH-DFT) at the quantum mechanical level. We selected representative arginine/methionine pairs from hydrophobic (n = 34) (Fig. 2e) and polar (n = 48) (Extended Data Figs. 7a & 7b) environments, and for each pair, we performed geometry optimization to converge to its closest local minimal-energy conformation prior to the energy calculations. For the DH-DFT calculations, we used methyl-guanidinium and dimethyl-sulfide to represent arginine and methionine, respectively (Fig. 2e). The methionine Sδ engages with the arginine’s guanidinium in a Van der Waals-like interaction that can contribute up to -3.3 kcal/mol in a low dielectric environment (Fig. 2e) as in the hydrophobic pocket of the LUJV spike (Fig. 2d) and up to -2.5 kcal/mol in a high dielectric environment (Extended Data Fig. 7a). Pairs with the more abundant angle between the arginine and methionine planes near 90°, as seen in the PDB-derived statistical data, also show somewhat stronger interactions (Fig. 2e). Taking into account the sulfur-guanidino interaction and the additional contribution of the aliphatic tails, the Arg130-Met221 interaction is a significant quaternary-enabled interaction for the NRP2/LUJV spike complex.

The dense array of N-linked glycans that viruses typically mount on their viral spike complexes helps them to conceal important functional sites, like the receptor binding site, for example, from the humoral immune system. In the case of LUJV, the apex of the spike is decorated by several N-linked glycans that are visible in the density maps (Fig. S1). Of these glycans, two are particularly close to NRP2: a glycan attached to Asn112 on the cognate GP1 domain and a glycan attached to another Asn112 on a neighboring domain (Fig. 2f). NRP2 fits very tightly between these two glycans. The Asn112-linked glycan on the cognate GP1 domain, as modeled in the EM structure, will clash with NRP2 unless it tilts (Fig. 2f). Indeed, the first GlcNAc of this glycan is visible in the crystal structure of NRP2/GP1, and it assumes a tilted orientation compared with the glycan in the EM structure (Fig. 2f).

Having these glycans at a contact distance from NRP2 may affect complex formation by either sterically restricting access of NRP2 to the spike or, on the other hand, promoting complex formation by contributing molecular interactions with NRP2. Interestingly, pseudo-viruses decorated with LUJV spikes that do not have glycans attached to Asn112 (*i.e.*, having a T114A mutation that abrogates the N-X-T/S glycosylation motif) can infect cells as efficiently, and perhaps even more efficiently, compared with pseudo-viruses equipped with the WT spikes (Fig. 2g). This observation suggests that the Asn112-linked glycans do not contribute for the binding of NRP2, and may even restrict binding by steric interference. Testing the capacity of NRP2-based competitors to prevent cell entry of pseudo-viruses further strengthens this notion: A bulky NRP2-Fc immunoadhesin, made of the first CUB domain of NRP2 fused to Fc portion of IgG1, seems to neutralize LUJV better when the Asn112-linked glycan is absent (Figs. 2g & Extended Data Fig. 8), and a smaller reagent that comprises only the first CUB domain of NRP2 (NRP2-His) achieves better IC_50_ values for neutralization compared with NRP2-Fc (Fig. 2g). Hence, the Asn112 glycan reduces the overall capacity of the LUJV to interact with its NRP2 receptor by imposing steric constraints.

### Shared and unique properties of the LUJV spike complex

Besides the structure of the LUJV spike complex, structural information for a complete arenaviral spike complex is, so far, available only for the LASV ^17^. Despite sharing a similar overall organization, superimposing the two structures yields an RMSD value of 22.5 Å for 1095 shared Cα atoms. Such a high RMSD value indicates different relative orientations of the GP1, GP2, and SSP domains. Indeed, the similarity of the two spikes is visually apparent (Fig. 3a), and superimposing the individual domains yields RMSD values of 8.2 Å, 3.0 Å, and 1.0 Å for the GP1 (155 Cα atoms), GP2 (176 Cα atoms), & SSP (33 Cα atoms), respectively (Fig. 3b). While the transmembrane region and the juxtapose GP2 region are structurally similar, the GP1 domains themselves and their relative orientation in the spikes greatly differ between LUJV and LASV (Figs. 3a & 3b). The mutual packing of the GP1 domains in LUJV seems to be looser compared with the packing in LASV (Fig. 3c); Three narrow cavities are forming between the three GP1 subunits of LUJV, and the interacting surfaces are limited to the vicinity of the spike’s symmetry axis (Fig. 3c). In LASV, no such cavities are formed, and the packing of the GP1 domains spans a larger interface (Fig. 3c). Nonetheless, in both spike complexes, critical elements for the trimerization are the C-terminal ends of the GP1s that contain the SKI-1 recognition sequences.

**Figure 3.**
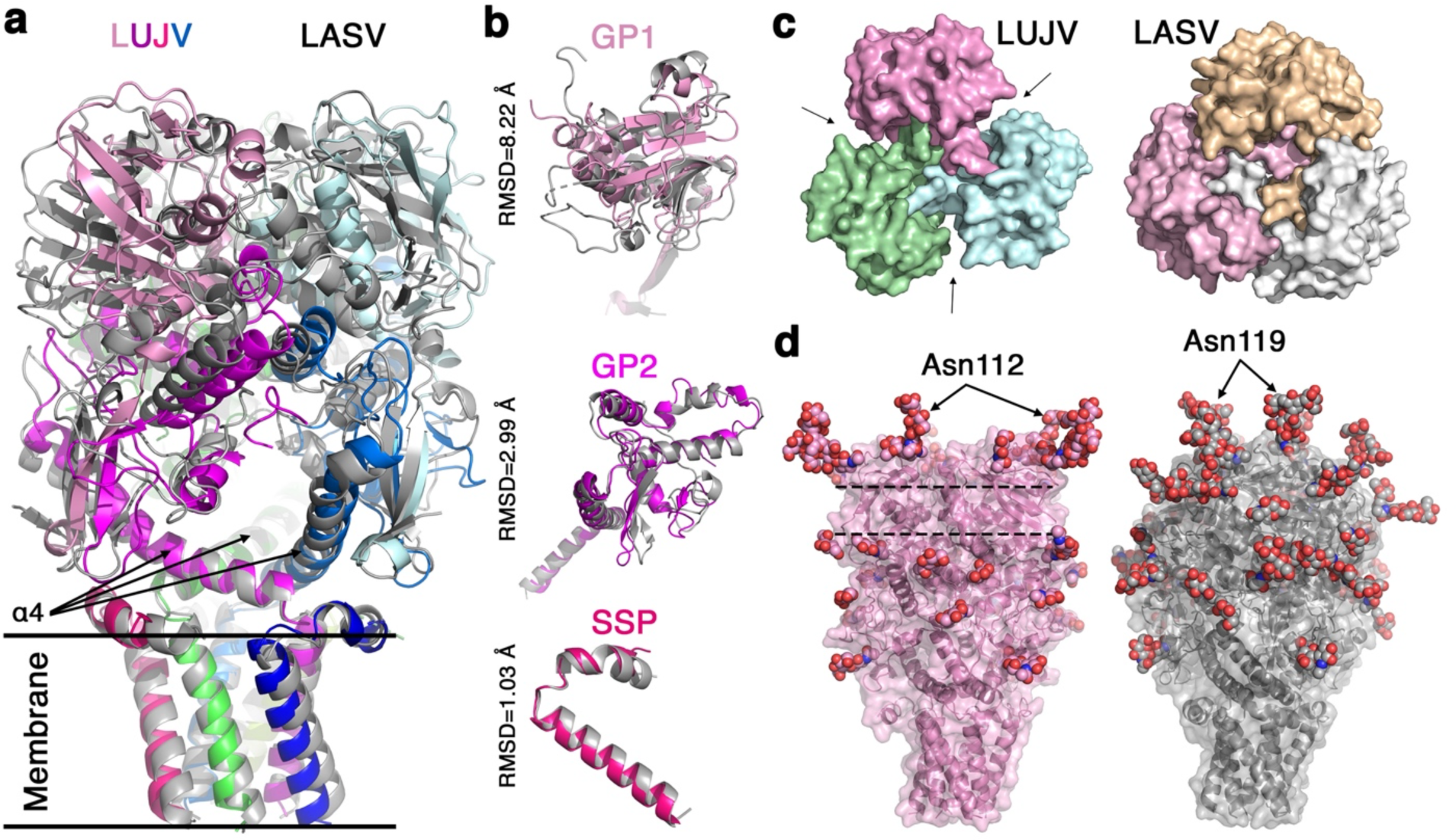
Comparison of the LUJV and LASV spike complexes. **a**. Superimposition of the complete LUJV and LASV (PDB ID: 7PUY) spikes, calculated based on the helix α4 regions of the GP2s. The LUJV spike is colored using the same coloring scheme as in Figure ‘1’, and the LASV spike is shown in grey. **b**. Superimpositions of the individual domains of LUJV and LASV. RMSD values for matching mutual Cα atoms are indicated. **c**. Top views of the GP1 domains from the LUJV (left) and LASV (right) spikes, using surface representations and a distinct color for each GP1 domain. Arrows point to gaps between the GP1 domains of the LUJV spike complex. **d**. Glycosylation of the spike complexes. N-linked glycans are shown as spheres on the LUJV (left) and the LASV (right) spikes. The glycan sites that are closest to the spikes’ three-fold symmetry axes are indicated. Two dashed lines highlight a region of the LUJV spike that has no glycans.

Arenaviruses strategically place glycans on their spikes to evade neutralizing antibodies and reduce clearance ^18^. LUJV binds its receptor such that NRP2 overlaps with the spike’s symmetry axis (Fig. 2b), and hence it must leave the apex of the spike devoid of glycans (Fig. 3d). The glycan closest to the symmetry axis is attached to Asn112 (Fig. 3d), which is at a contact distance from NRP2 (Fig. 2f). Since the matriglycan receptor of LASV fits into a small pocket at the apex of the spike ^17^, the LASV spike can have glycans that are much closer to the three-fold symmetry axis compared with the LUJV spike. Indeed, the Asn119-linked glycan of LASV is apparently closer to the apex than the Asn112-linked glycan of LUJV (Fig. 3d). This difference makes the receptor binding site of LUJV more exposed to antibodies compared to LASV. Interestingly, while the N-linked glycans on LASV seem to be evenly distributed on the surface of the spike, there is an evident belt on the surface of the LUJV spike complex that is completely lacking N-linked glycans (Fig. 3d). While viral glycan shields are never completely sealed, and will always leave patches that are free of glycans, as in the case of LASV (Fig. 3d) or other viral class-I spike complexes (Extended Data Fig. 9), having a complete belt without glycans seems to be unusual. Having such a belt free of glycans could be a pure coincidence, or it may confer some beneficial properties to the spike, like creating an antibodies-accessible immunodominant region to subvert immune response.

### Evaluating the potential of an anti-LUJV immunoadhesin

Previously, we demonstrated that host-derived receptors could be utilized to make powerful immunoadhesins against zoonotic viruses ^19,20^. In the case of LUJV, however, the exact host is unknown, and the NRP2 interface that GP1 recognizes is highly conserved in mammals ^16^, complicating the construction of high-affinity immunoadhesin. Potentially, NRP2-Fc may be used as an immunoadhesin (Fig. 2g), but such a reagent is likely to have insufficient potency for clinical use. As we noted above, the N-linked glycan on Asn112 restricts the binding of the LUJV spike to NRP2 (Fig. 2f & 2g). A close inspection of the LUJV spike’s structure with NRP2 superimposed reveals that the N-linked glycan on Asn112 is clashing with Tyr128 of NRP2 (Fig. 4a). Eliminating the bulky side-chain of Tyr128 in NRP2 with a Y128G mutation increases the potency of NRP2-Fc by nearly tenfold, compared with the WT NRP2-Fc (Fig. 4b), as measured using MLV-based pseudotyped viruses bearing the LUJV spike complex. Moreover, the structure suggests that Gln47 of NRP2, in its favorite rotamer, would be forced to assume a different rotamer to avoid clashing with a neighboring GP1 domain (Fig. 4a). Restricting the available rotamers for Gln47 confers an entropic cost for binding and hence reduces affinity. Indeed, eliminating the Gln47 side chain with a Q47G mutation improves the neutralization of NRP2-Fc (Fig. 4b). Lastly, the structure further suggests that Ile108 will be in very close proximity, perhaps will even clash, with Asn112-linked glycan from the neighboring GP1 monomer (Fig. 4b). Introducing an I108S mutation to NRP2-Fc also results with a more potent neutralization of LUJV (Fig. 4b).

**Figure 4.**
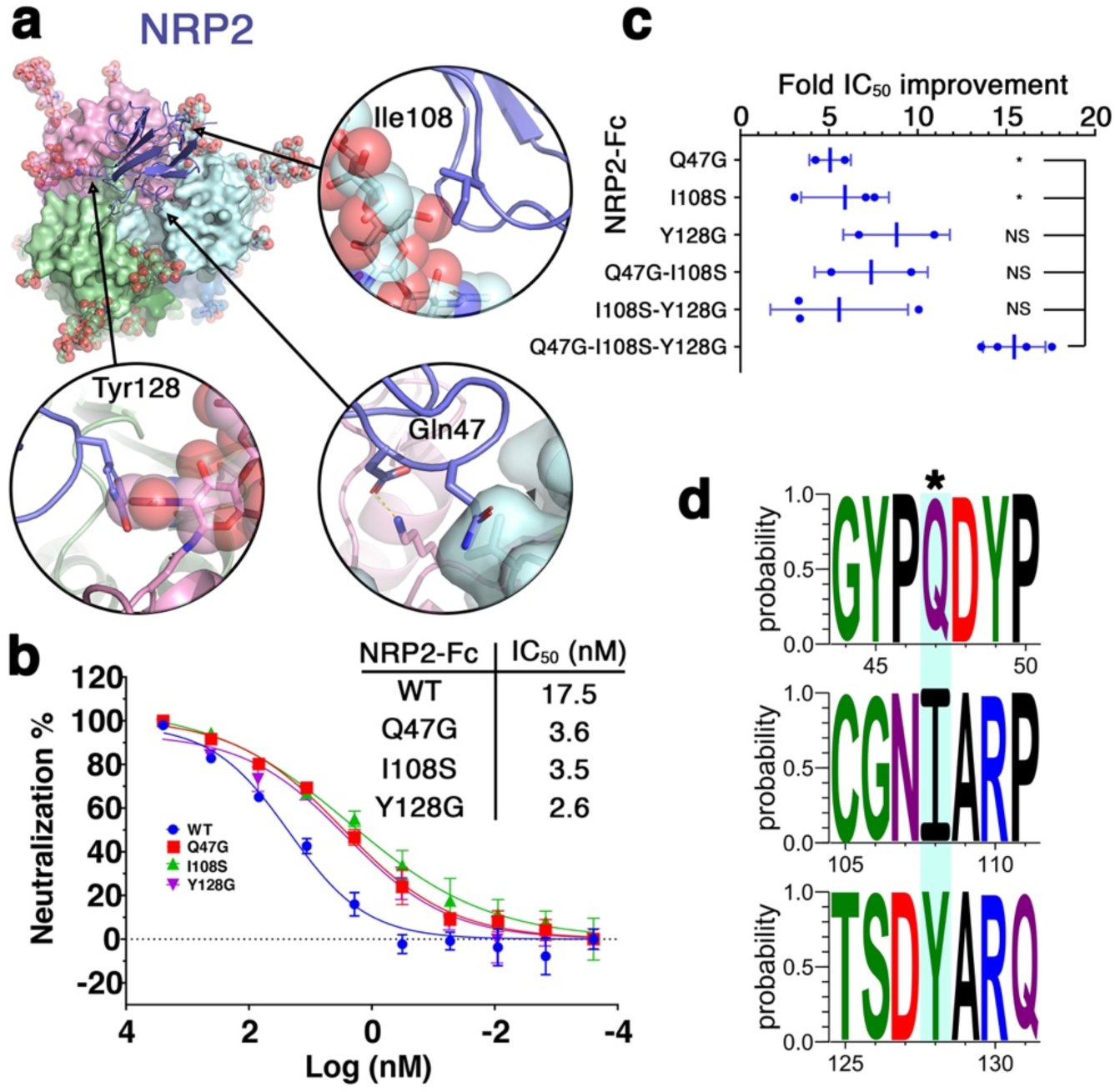
Mutations enhance neutralization of LUJV by NRP2-Fc. **a**. Closeup views of the three residues that are predicted to interfere with the binding of NRP2 by the LUJV spike. Tyr128 of NRP2 is on the cognate GP1 (pink), and Ile108 and Gln47 are on adjacent GP1 (light blue). **b**. Neutralization of LUJV pseudoviruses by NRP2-Fc WT and the indicated mutants. The graph is from a representative experiment (each data point is the average value from four technical repeats, and error bars show standard deviations), and the IC_50_ values are averaged values from several independent experiments. **c**. Fold improvement of neutralization IC_50_ values of the NRP2-Fc mutants compared with WT NRP2-Fc. Each point represents an independent experiment. Averaged values are indicated, and error bars show standard deviations. Statistical significance for the differences between the triple mutant (Q47G-I108S-Y128G) and the single or double mutants are indicated (NS - not significant, * - p<0.05 two-tailed Student’s T-test). **d**. Conservation of the mutated sites. Multiple sequence alignments of 447 NRP2 sequences from mammals are summarized as WebLogos around the Gln47, Ile108, and Tyr128 sites (top, middle, and bottom, respectively). In each WebLogo, the middle residue (noted with an asterisk) is the mutation site.

We next tried to combine two or three mutations in a single NRP2-Fc reagent. Combining two mutations did not have any added value compared with the single mutations, reaching 5-10-fold improvements in IC_50_ values compared with WT NRP2-Fc (Fig. 4c & Extended Data Fig. 10). Combining all three alterations resulted in a more potent reagent that shows up to 15-fold improvement compared to WT (Fig. 4c). The combination of the three mutations seems to improve the capacity of NRP2 to engage with and neutralize the LUJV spike complex, albite this improvement is not statistically significant compared to some of the other combinations (Fig. 4c), likely due to the limited sample size. Nevertheless, NRP2^(Q47G, I108S, Y128G)^-Fc achieves an improved IC_50_ value of 1.6 nM (*i.e.*, 0.12 μg/ml) compared with NRP2-Fc, an improvement that is statistically significant (p<0.0002, Student’s two-tailed T-test). Interestingly, this NRP2-Fc variant achieves fairly high neutralizing capacity despite the fact that its binding mechanism clearly lacks avidity (Extended Data Fig. 10). The absence of avidity is further noted by the shallow slopes (Hill slopes smaller than 1) of the neutralization curves (Fig. 4b, Extended Data Fig. 10, & Extended Data Fig. 8). As noted above, only a single NRP2 can bind to the LUJV spike at any given time (Fig. 2b). Hence, avidity could only be enabled if the NRP2-Fc is able to bind two adjacent spikes. We, therefore, postulated that the hinge connecting the NRP2 CUB domain to the Fc is not sufficiently long to allow such binding. Extending the hinge with an additional 10 residues-long GS-linker, however, did not improve neutralization (data not shown), indicating that avidity was not enabled.

## Discussion

Being the second arenaviral spike complex whose complete structure has been determined so far, the single particle cryo-EM structure of the LUJV spike sheds light on common structural elements of arenaviral spikes. Both LUJV and LASV spikes organize as a six-helical bundle in the membrane, where the SSP functions as a stabilizing element, holding the three central GP2 transmembrane helices wound around each other. The observed topology of SSP in the membrane in both spikes is the same, indicating that topology rearrangement is common for arenaviral spikes. Also, the C’-end of GP1, which results from the cleavage by SKI-1, is an important trimerization factor at the apex of both spikes. Albeit the actual structures differ, the critical function of the SKI-1 recognition motif in mediating trimerization is preserved. This special structural role adds to the primary function of this motif in cleaving the GPC polypeptide chain for the spikes’ maturation. This structural role further rationalizes why these arenaviruses adhere to SKI-1 as their cellular protease for maturation rather than evolving to utilize other cellular proteases like furin, for example, which other viruses use to enhance their virulence ^21^.

Another important theme that emerges from this study is the stabilizing effect on the viral spike complex that is predicted to result from receptor binding. The quaternary-enabled interactions that NRP2 forms with the spike (Fig. 2c, 2d, & Extended Data Fig. 6) inevitably favor the trimeric form of the spike by locking the C’-ends of the GP1 subunits in their bound conformation. These quaternary interactions include the identified interaction between Met221 and Arg130. Such interaction between sulfur atoms of methionine and the guanidino groups of arginine is a previously overlooked molecular attribute that contributes to the structural stability of proteins. Receptor binding also stabilizes the spike in the case of LASV. Similarly to LUJV, the C’-ends of the GP1 subunits in the spike of LASV are also critical elements in the formation of the matriglycan binding site ^17^. Likewise, binding to the matriglycan receptor favors the trimeric form of the LASV spike. In addition to stabilizing the trimeric form, in the case of the LASV spike, matriglycan also serves as an N’-cap for exposed helices in the GP1 subunits. Therefore, the binding of matriglycan further stabilizes the folded state of the GP1 domains. Interestingly, both LUJV and LASV are known to dissociate from their primary cell-host receptors in a pH-driven process and utilize a secondary receptor for triggering ^4,22,23^. Taken together, the pH-driven dissociation from NRP2 in the case of LUJV or matriglycan in the case of LASV also destabilizes the trimeric organization of the spikes. This may be a required step on the path to the induction of membrane fusion.

The sub-optimal binding of the LUJV spike to human NRP2, as exemplified by the identification of three different binding-enhancing alterations to NRP2, is intriguing. NRP2 is a conserved protein, and analysis of 447 NRP2 sequences from mammals (Extended Data Appendix 2) indicates that Gln47, Ile108, and Tyr128 are completely conserved (Fig. 4d). Viruses typically adapt to their natural host receptor ^24^, and the observed sub-optimal compatibility with the human NRP2 could be explained in two different ways: First, it might be that the natural host of LUJV is not a mammal, and has a cell surface receptor with a more compatible sequence than the human NRP2. Second, it might be that LUJV sacrifices some binding affinity to its receptor in order to sterically conceal its receptor-binding site from the immune system. Regardless of the exact reason for this sub-optimal binding, its mere existence provided an opportunity to construct an NRP2-based immunoadhesin that binds the LUJV spike complex better than the human NRP2. While in the absence of avidity, the potency of the NRP2-Fc immunoadhesin could be insufficient for clinical use, the NRP2^(Q47G, I108S, Y128G)^ variant itself is a promising reagent, warranting further research since having an off-the-shelf remedy for a future LUJV outbreak is highly desired.

## Data availability

Coordinates file and experimental density maps were deposited at the PDB and EMDB under accession codes 8P4T and EMD-17428, respectively.

## Acknowledgments

The Diskin lab is supported by research grants from the Ernst I Ascher foundation, Ben B. and Joyce E. Eisenberg Foundation, Estate of Emile Mimran, Jeanne and Joseph Nissim Center for Life Sciences Research, Dov and Ziva Rabinovich Endowed Fund for Structural Biology, Donald Rivin, Stanley and Tanya Rossby Endowment Fund, Natan Sharansky, Dr. Barry Sherman Institute for Medicinal Chemistry, as well as from the Israel Science Foundation (grant No. 209/20).

## Author contributions

R.D. conceived and oversaw this research. M.E-A. produced and purified proteins. M.E-A.& R.D. performed EM analysis and solved the structure. M.E-A. & S.B. performed infectivity and neutralization assays. R.S. performed a statistical analysis of structural information from the PDB. R.C. & K.L. performed DH-DFT calculations and analyzed data. H.C-D. provided reagents. All authors contributed to the data analysis. R.D. & M.E-A. prepared the manuscript with the help of all the other authors.

## Methods

### LUJV GPC production and purification

Expression of the full-length LUJV GPC was carried out in HEK 293F cells (Invitrogen) using FreeStyle Medium (Life Technologies). Cells were grown to a density of approximately 1.0 × 10^6^ cells per ml before transfection. HEK 293F cells were transfected using 40 kDa polyethylenamine (PEI-MAX) (Polysciences) at 1 mg ml^−1^, pH 7 with DNA at a ratio of 1:2 (DNA:PEI solution). Codon-optimized LUJV GPC was chemically synthesized (Genescript) and then subcloned with a C-terminal Flag tag into pcDNA3.1 using BamHI–NotI restriction sites. The LUJV GPC-expressing cells were collected at 48 h post-transfection by centrifugation at 700xg, 4 °C for 5 min. Membranes were then resuspended in a cold lysis buffer (10 mM Tris, 150 mM NaCl, 100 μM MgCl_2_, 1 mM EDTA, 100 μM phenylmethyl sulfonyl fluoride (PMSF), 15% glycerol) and homogenized for 5 min on ice. The lysis mixture was then incubated while rotating for 1 h at 4 °C. A second homogenization was carried out and the lysis mixture was centrifuged at 33,300xg for 25 min, 4 °C. The supernatant was discarded, and pellets were dissolved in solubilization buffer (20 mM Tris, 150 NaCl, 15% glycerol, 3% (% w/v) n-dodecyl-β-D-maltoside (DDM; Anatrace)). The solubilization mixture was then homogenized and incubated for 4 h; after which it was centrifuged at 370,000xg for 25 min, 4 °C. The supernatant of this solubilization step was then incubated overnight with 50 μl of EZview red anti-Flag beads (Sigma Aldrich). The insoluble material from the solubilization step was discarded. The anti-Flag beads were then spun down (800xg, 2 min) and washed by subsequently decreasing amounts of glycerol to 0.75% and DDM to 0.03%. The protein was eluted after the last washing step with 100 μl of 0.40 mg/ml of 1× Flag peptide (Genescript) in a buffer containing 0.03% DDM, 20 mM Tris-HCl, 150 mM NaCl, from a 1-hour incubation on ice. For western blot analysis, anti-Flag primary antibody (Thermo Fisher) was used at a 1:1,000 dilution with horseradish peroxidase (HRP)-conjugated anti-mouse secondary antibody (Thermo Fisher) at a 1:10,000 dilution.

### Cryo-EM image acquisition, data analysis, and 3D reconstruction

A purified LUJV spike sample (3.5 μl) was applied on glow-discharged (6 s, 12 mA; Pelco easiGlow, Ted Pella) graphene oxide Quantifoil copper grids, R1.2/1.3, (Electron Microscopy Sciences) using a Vitrobot system (Thermo Fischer/FEI) (4.5 s blotting time, 4 °C, 100% humidity). Samples were incubated on the grid for 1 min before blotting was carried out. cryo-EM data was then collected on the Titan Krios microscope (FEI) operated at 300 kV, using a Gatan K3 direct detection camera. The beam size was 705 nm diameter (fringeless illumination), the exposure rate was 18 e^−^ s^−1^ pixel^−1^, and movies were then obtained at 105,000× magnification with a pixel size of 0.793 Å. The nominal defocus range was −0.6 to −1.8 μm. A total of 11,381 movies were automatically collected using EPU. Data processing was carried out with the cryoSPARC v4.1.2 suite. Patch motion correction and patch CTF estimation were carried out using cryoSPARC Live. Blob picking was used to pick 5,877,899 particles. Particles were extracted using a 256-pixel box, and the data set was cleaned from junk particles by carrying iterative 2D classifications, followed by 3D classifications. The final map was obtained from non-uniform refinement, imposing C3 symmetry from 87,868 particles. This map was then filtered using DeepEMhancer ^25^ to obtain the working map.

### Model building, refinement, and analysis

An initial model was generated by docking the GP1 structure (6GH8) and the GP2 structure (7PUY) into the density map using Chimera. Then, using Coot and real-space refinement in Phenix, we manually completed and refined the model of the spike.

### Pseudoviral Particle Production

MLV virus-like particles (VLPs) pseudotyped with LUJV GPC were produced by transfecting retroviral transfer vector pLXIN-Luc encoding luciferase as a reporter gene together with LUJV-flag or mutated LUJV-flag in pcDNA3.1 into the GP2-293 retroviral packaging cell line (Clontech). GP2-293 cells were seeded at 5×10^6^ on 10-cm plates and transfected 24 h later with 5 µg of LUJV-flag/LUJV-flag-mut and 5 µg Luciferase using Lipofectamine 2000 (Invitrogen). Cells’ media were replaced 5 h later to full medium, i.e., DMEM (Biological Industries) supplemented with 1% Pen-Strep (v/v), 1% Glutamine (v/v), and 1% sodium pyruvate (v/v). At 48 h post-transfection, media containing pseudoviruses were harvested, and VLPs were concentrated 10 times by the addition of PBS/8 % (w/v) PEG 6000 (Sigma), incubation at 4° C for 24 h, centrifugation at 10,000xg for 20 min and resuspension in full medium. The concentrated VLPs were stored at -80° C until use. Mutated genes of LUJV-flag were generated by PCR mutagenesis.

### Infectivity Assays

For infectivity assays, HEK293T cells were seeded on a poly-L-lysine-precoated white, chimney 96-well plate (Greiner Bio-One). Cells were left to adhere for 3 h, followed by the addition of LUJV VLPs. Cells were washed from the viruses at 18 h post-infection, and luminescence from the activity of luciferase was measured at 48 h post-infection using a Tecan Infinite M200 Pro plate reader after applying Bright-Glo reagent (Promega) to the cells.

### NRP2-his and NRP2-Fc Purification

Expression constructs of the first CUB domain of Neuropilin-2 (NRP2) fused to a 6 x His tag or Fc were previously constructed ^16^. Expressions of the proteins were carried out in HEK 293F cells (Invitrogen) using FreeStyle Medium (Life Technologies). Cells were grown to a density of approximately 1.0 × 10^6^ cells per ml before transfection. HEK 293F cells were transfected using 40 kDa polyethylenamine (PEI-MAX) (Polysciences) at 1 mg ml^−1^, with DNA at a ratio of 1:3 (DNA:PEI solution). Media containing the NRP2 constructs were collected at 6 d post-transfection. The NRP2-his was purified using a HiTrap IMAC FF Ni^2+^ (GE Healthcare) affinity column and Superdex 75 10/300 size exclusion chromatography (GE Healthcare) and then concentrated to an optical density at 280 nm (OD_280_) of 4, in 20 mM Tris-HCl pH 8.0, 150 mM sodium chloride, 10 µM CaCl_2_, 0.02% (wt/vol) sodium azide using an Amicon concentrator (Millipore). The NRP2-Fc was purified using a Protein-A affinity column (GE Healthcare). NRP2-Fc point mutations were introduced by PCR mutagenesis using Kapa HiFi DNA polymerase (Kapa Biosystems), according to the QuikChange site-directed mutagenesis manual. Mutated variants were expressed in HEK 293F cells. Media were collected 6 d post-transfection. The mutated proteins were purified from the media using a Protein-A affinity column (GE Healthcare).

### Bioinformatic structural analysis

A total of 34,376 structures from the PDB that were determined at 2.2 Å resolution or better and are unique (sequence similarity cutoff of 90%) were used for our analysis. We employed a custom-made Python script (see Extended Data Appendix 1) to identify interacting arginine and methionine residues and further used this script to perform geometry analysis. Raw output from this analysis is included as supplementary data. Plots were generated using R.

### Calculating binding energies

The binding energies, which represent the energy difference between the optimized pairwise conformation and the energies of the optimized separated residues, were computed using the ORCA 5.0.3 software ^26^. These energies were calculated at the quantum mechanics (QM) level, employing the double-hybrid revDSD-PBEP86-D4 functional ^27^, with def2-qzvppd’ basis set, following a brief optimization at the revDSD-PBEP86-D4/QZ QM level. The choice of the DH-DFT functional revDSD-PBEP86-D4 ^27^ was influenced by its significantly accurate performance with both π-π and cation-π datasets ^28,29^. To save computational cost, arginine was represented as methylguanidinium, while methionine was represented as dimethyl sulfide. To account for structures located in hydrophobic regions, an implicit solvent model Conductor-like Polarizable Continuum Model (CPCM), was utilized with a dielectric constant of 4.0. In contrast, conformations in well-dissolved regions were calculated using CPCM (water) with a dielectric constant of 80.4. The partial charge was calculated using the CHELPG calculation scheme for the global minimum energy ^30^.

### Statistical analysis

Statistical analysis and determination of IC_50_ values were performed using Prism (GraphPad) software version 8.0.2. Student’s two-tailed T-test was employed for comparing groups. IC_50_ values were calculated by fitting the dose-response neutralization curves to a sigmoidal dose-response (variable slope) model using nonlinear regression analysis within Prism software. These IC_50_ values signify the concentration at which 50% neutralization of LUJV by the NRP2 variant was achieved.

### Bioinformatic sequence analysis

447 mammalian NRP2 amino acid sequences were aligned using the EMBL-EBI Clustal-Omega multiple sequence alignment tool ^31^. The Uniprot accession codes corresponding to these sequences can be found in Appendix 2. Sequence logos were generated by using WebLogo 3.1 (http://weblogo.threeplusone.com/).

