## Supplementary Information for "The Structure of the Lujo Virus Spike Complex"

For

This extended data includes:

Extended data Figures 1-10
Extended data Table 1
Appendixes 1 & 2

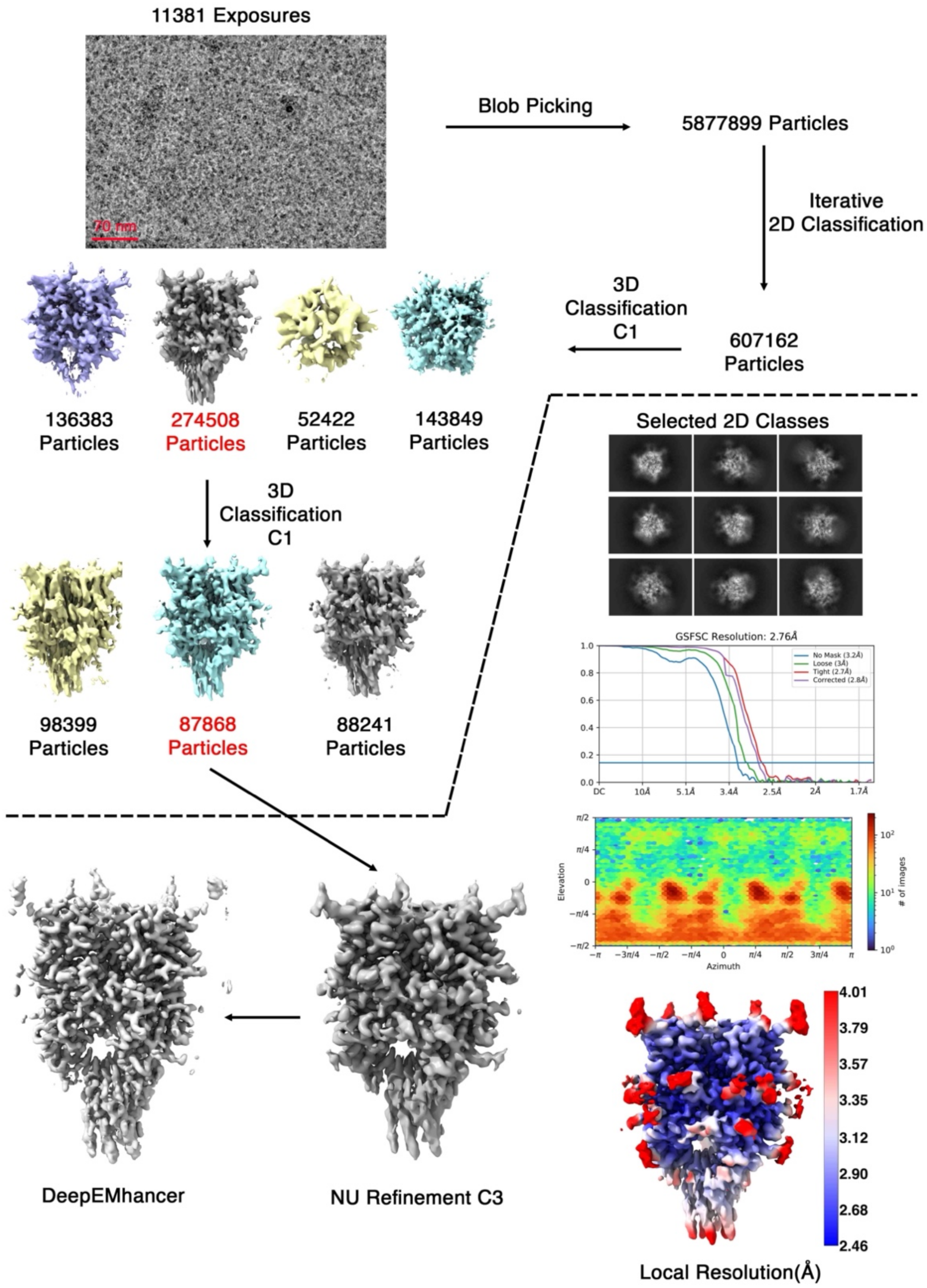


**Extended Data Fig. 1** | The reconstruction process of the electron microscopy density map. The procedure is schematically presented, starting with the upper-left image of a raw micrograph. The final map, its FSC curve, orientational distribution, local resolution, and a sharpened working map are shown.

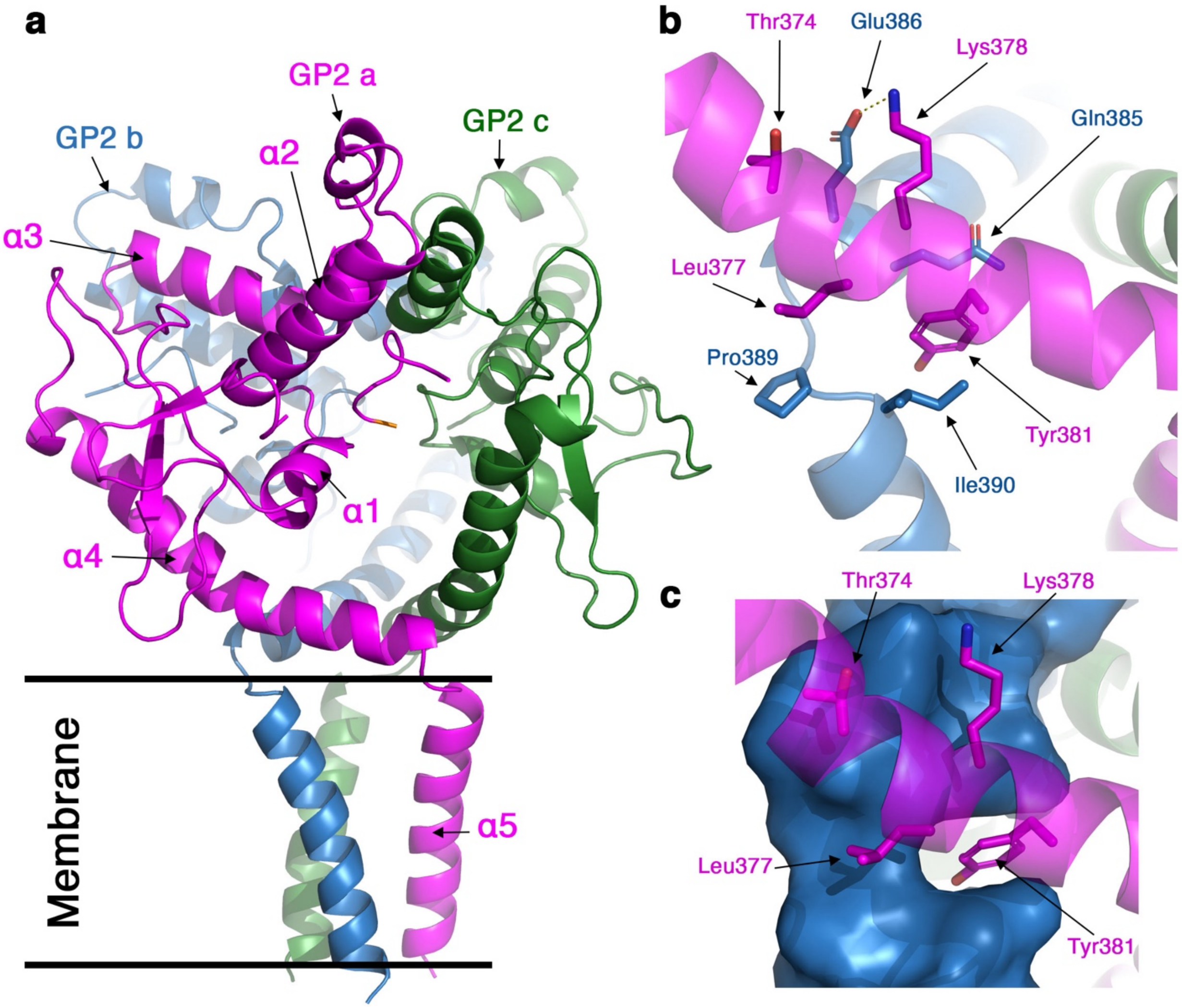


**Extended Data Fig. 2** | The interaction architecture between GP2 domains. **a**. Ribbon representation showing the three GP2 domains. The trans-membrane region is indicated. **b**. A close-up view of the interaction between $\alpha 4$ helices. **c**. Same as 'b' with one of the helices shown with a surface representation.

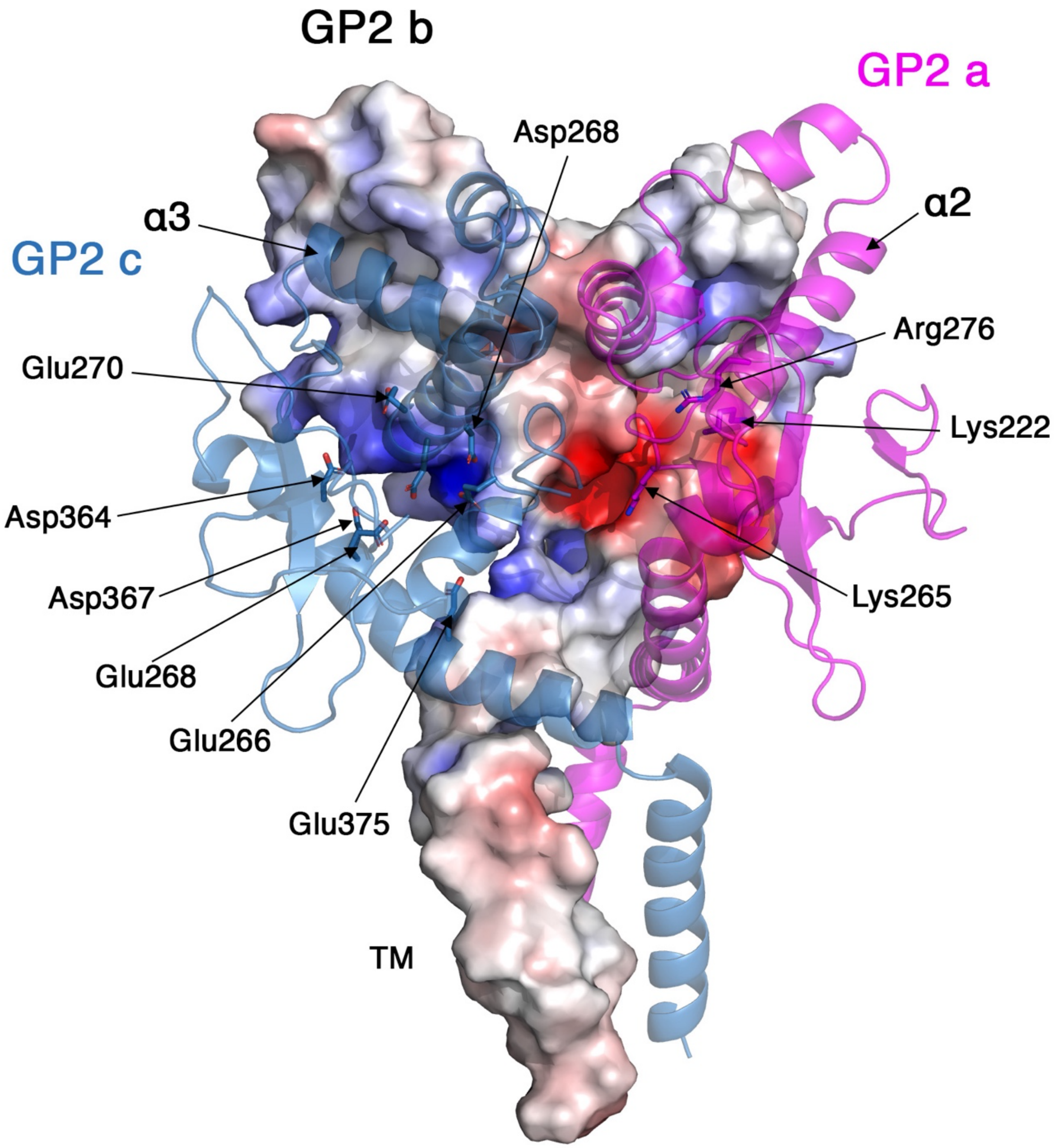


**Extended Data Fig. 3** | The electrostatic interaction between GP2 subunits. Three GP2 subunits are shown using a ribbon representation and a surface representation colored by electrostatic potential (red -5 kT/e; blue 5 kT/e).

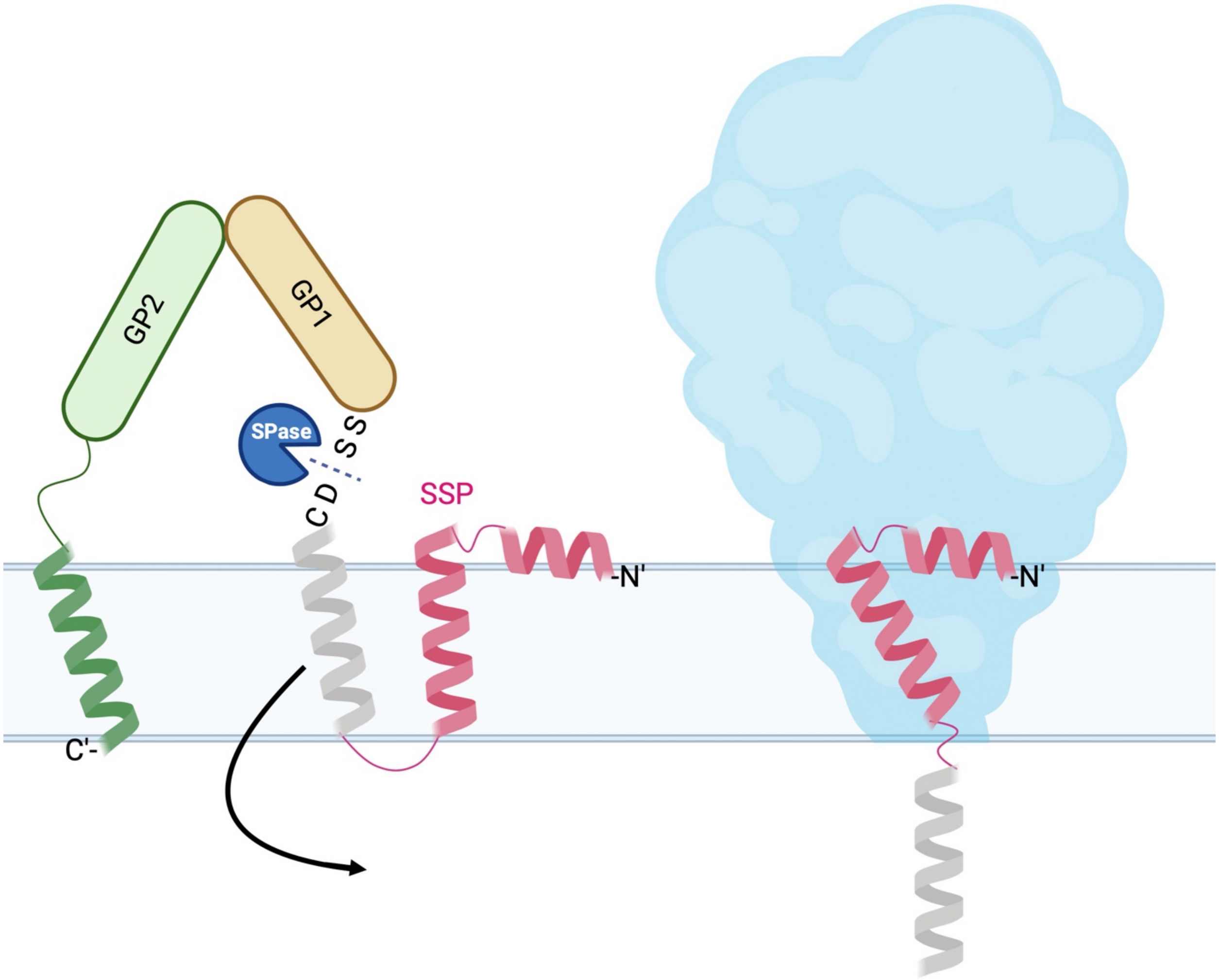


**Extended Data Fig. 4** | A schematic diagram showing the topology of the SSP before (left) and after (right) cleavage by SPase, in the context of the mature spike. The grey helix of SSP is not visible in the density map.

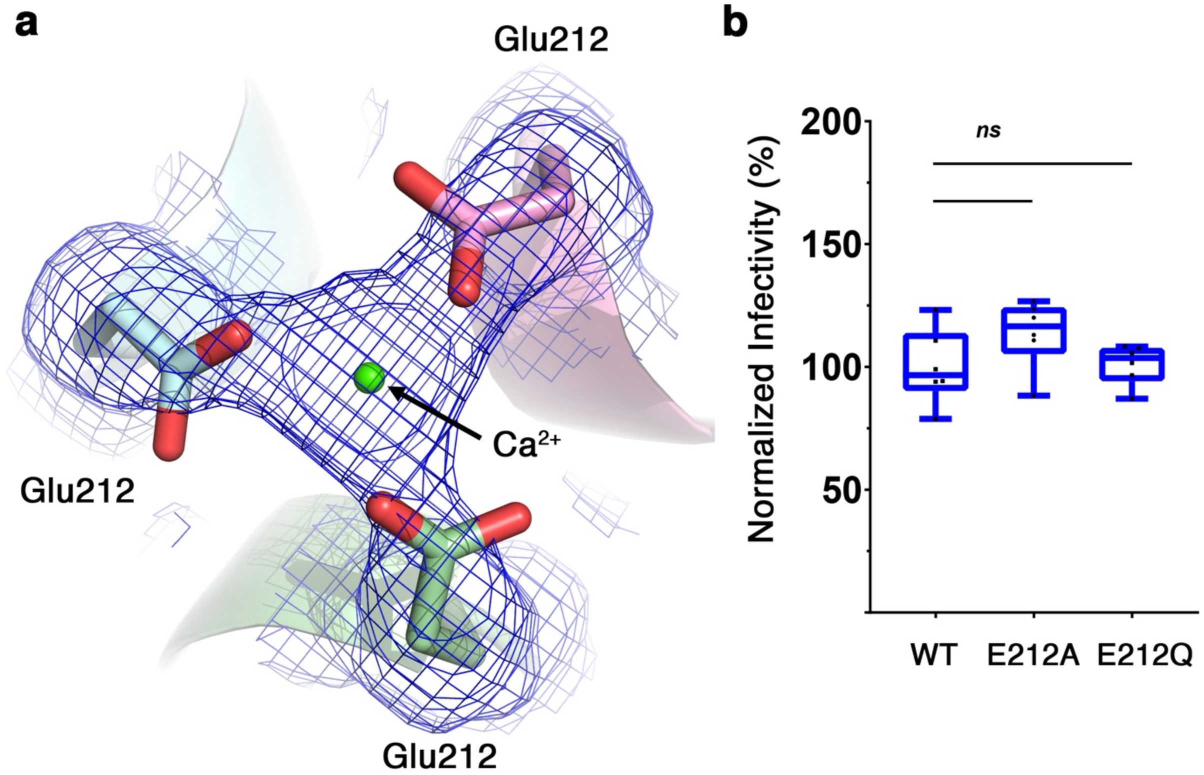


**Extended Data Fig. 5** | a. Density at σ=2 showing the Glu212 residues pointing toward the three-fold symmetry axis. The most likely metal ion is $Ca^{2+}$ which was included in the final model. b. Normalized infectivity of pseudo-viruses bearing either the WT or the indicated mutants at position 212.

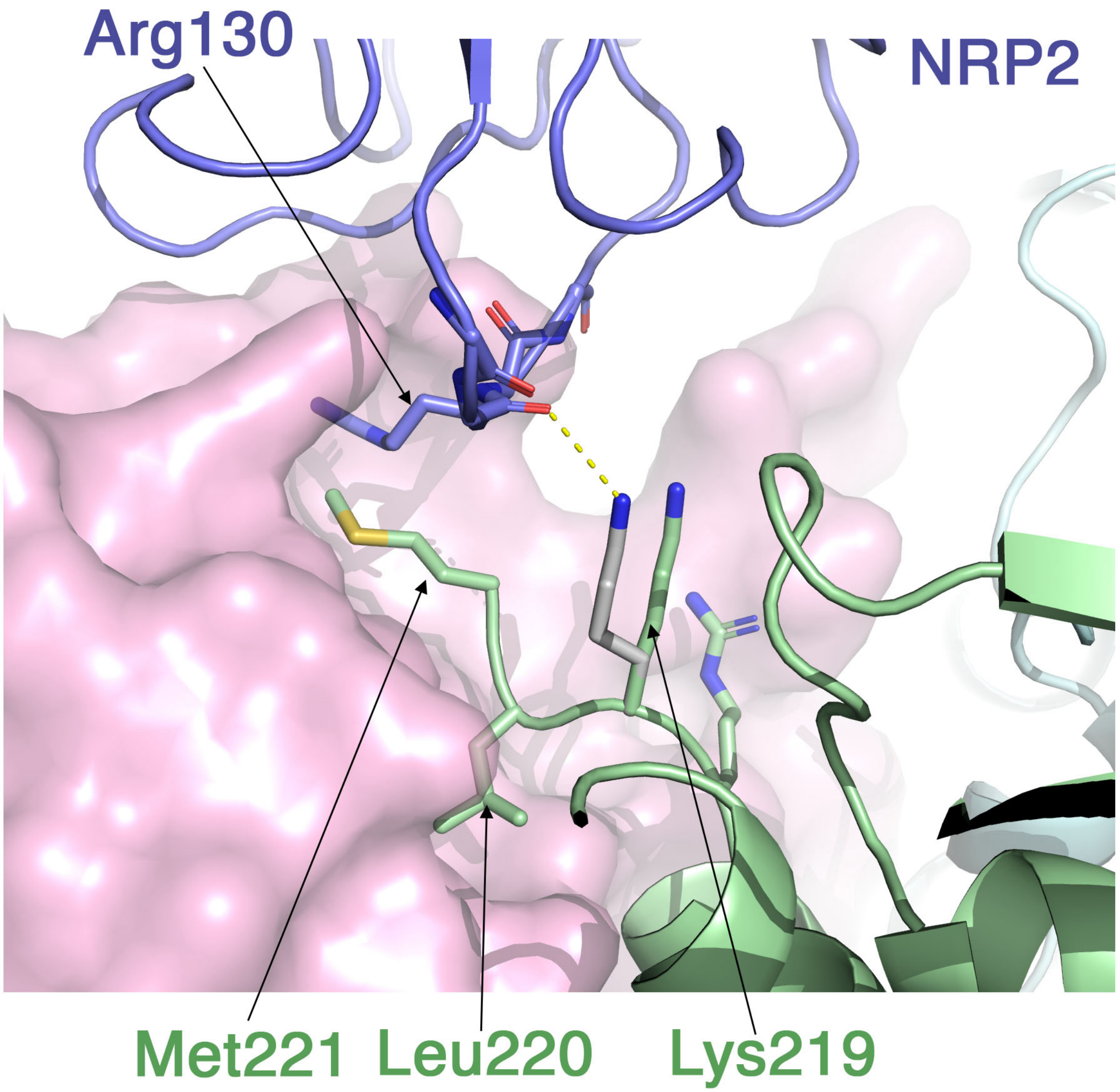


**Extended Data Fig. 6** | Potential salt-bridge between Lys219 and NRP2. The NRP2/GP1 pair (purple and pink, respectively) are shown with the neighboring GP1 (green). On top of forming a hydrophobic lid to Met221, the main-chain carbonyl of Arg130 can interact with Lys219, following a rotameric rearrangement. The observed rotamer in the EM structure is shown in green, and an alternative possible rotamer of Lys219 is shown in grey.

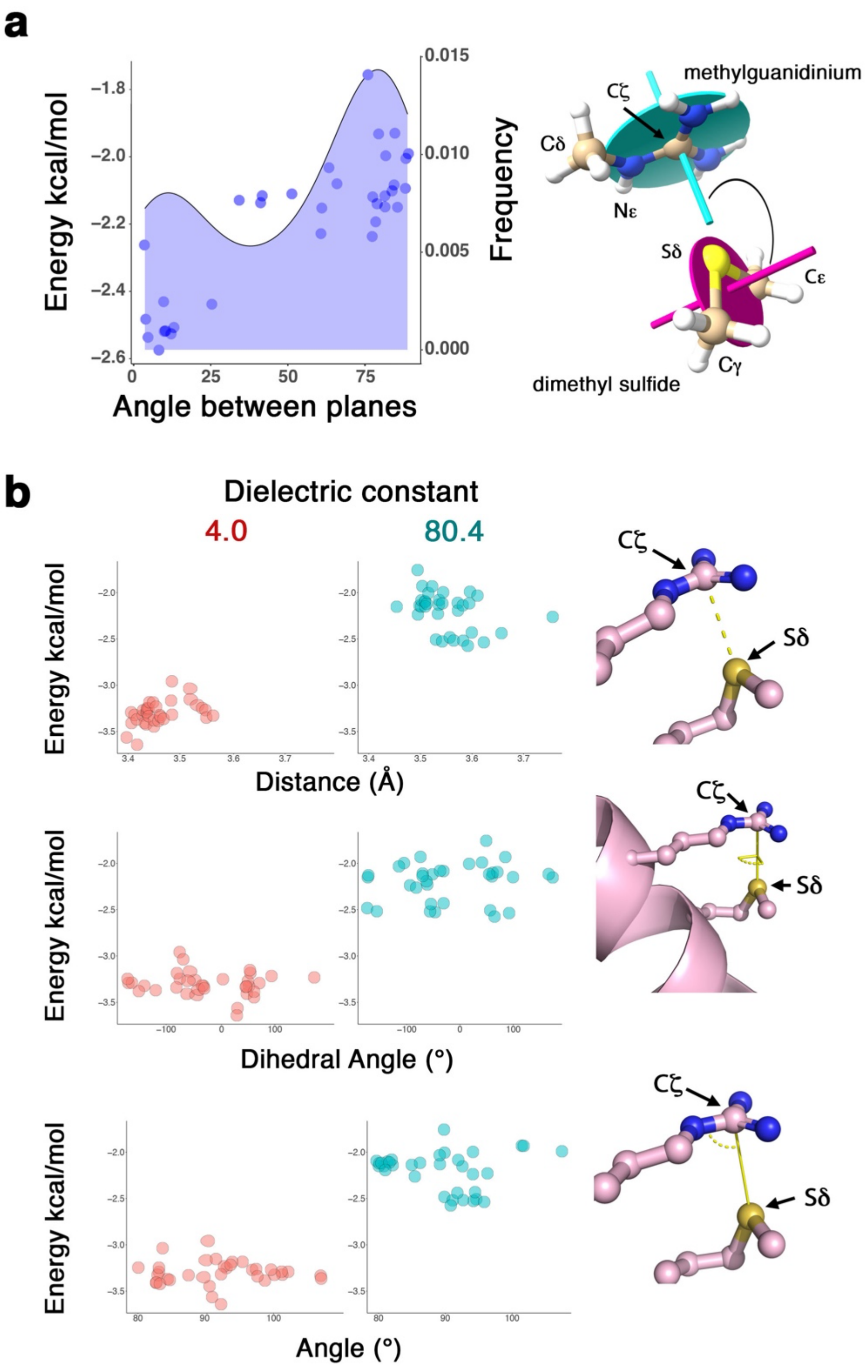


**Extended Data Fig. 7** | The sulfur – guanidinium interaction of methionine and arginine. **a**. the database-derived statistical frequency (blue chart) and the DH-DFT calculated binding energies (clue dots) for the interaction in a polar environment. **b**. distributions of geometrical attributes, and binding energies for the calculated arginine-methionine pairs, at high (blue) and low (red) dielectric environments.

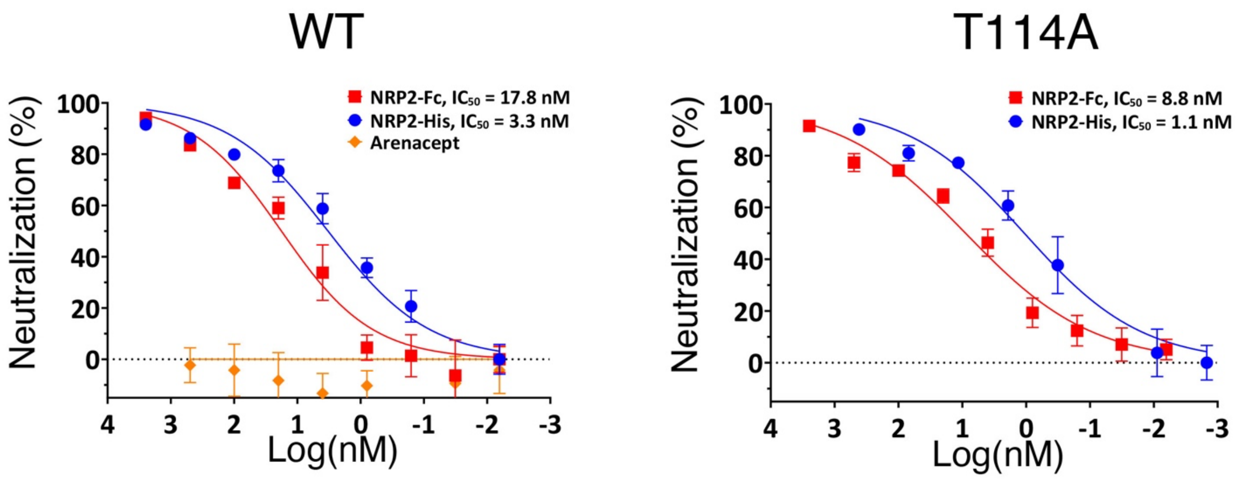


**Extended Data Fig. 8** | Neutralization of LUJV pseudo-viruses by NRP2 competing reagents. Neutralization of pseudo-viruses bearing either the WT (left) or the T114A (right) LUJV spike complexes. Neutralization by Arenacept that neutralizes NW arenaviruses [1] is used as a negative control. Error bars show standard deviations of n=6 technical repeats. These are representative experiments of three independent repeats.

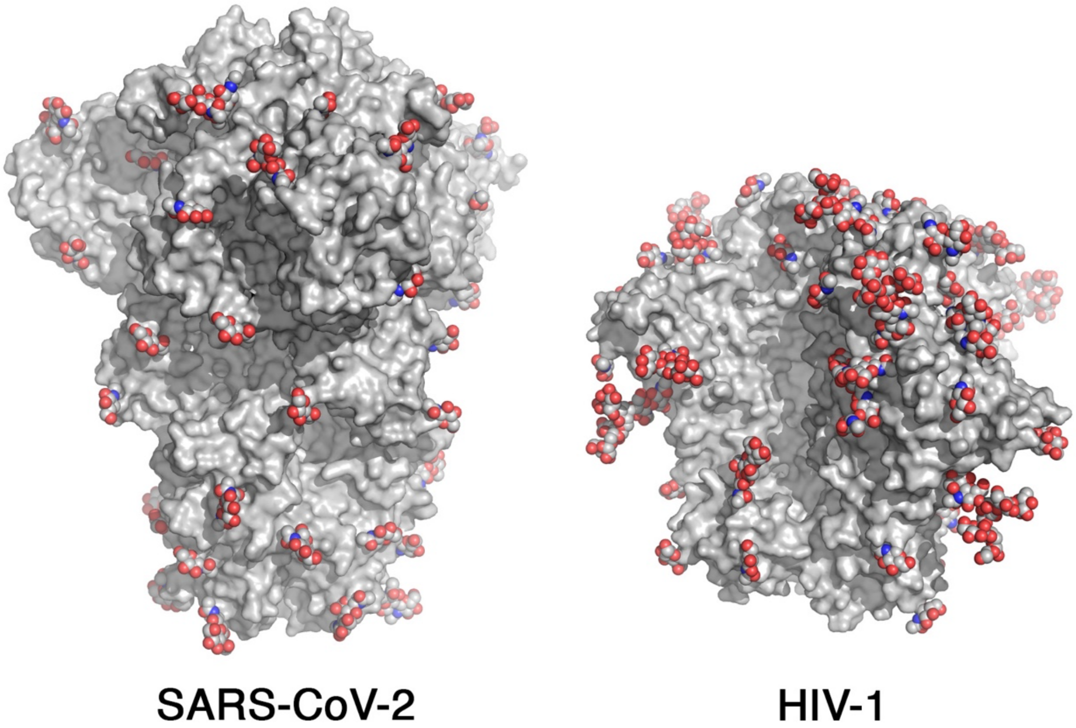


**Extended Data Fig. 9 Fig. S HIV** | Glycan shields on trimeric viral class-I spikes. The spike complexes of SARS-CoV-2 (PDB: 8DLW, left) and of HIV-1 (PDB: 7KDE, right) are shown in grey surface representations. N-linked glycans are shown as spheres.

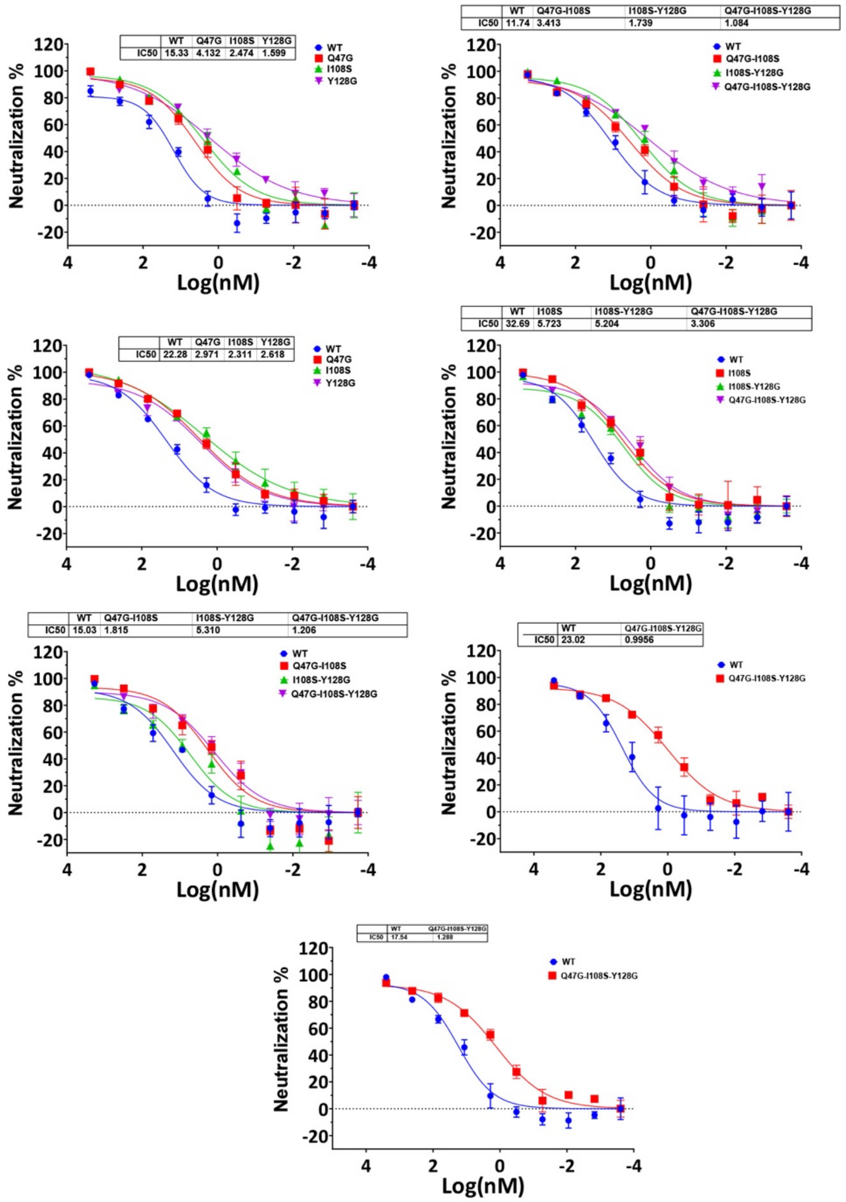


**Extended Data Fig. 10** | Neutralization curves of LUJV pseudotyped viruses by the indicated NRP2-Fc variants. Each graph summarizes the results from a single independent experiment. Calculated $IC_{50}$ values are indicated.

### Extended Data Table 1

| Model | PDB ID 8P4T | |
|---|---|---|
| **Composition (#)** | | |
| Chains | 16 | |
| Atoms | 9976 (Hydrogens: 0) | |
| Residues | Protein: 1122 Nucleotide: 0 | |
| Water | 0 | |
| Ligands | BMA: 6 | |
| | NAG: 39 | |
| | FUC: 3 | |
| | CA: 1 | |
| | MAN: 15 | |
| **Bonds (RMSD)** | | |
| Length (Å) (# > 4σ) | 0.005 (0) | |
| Angles (°) (# > 4σ) | 0.679 (0) | |
| MolProbity score | 1.72 | |
| Clash score | 16.49 | |
| **Ramachandran plot (%)** | | |
| Outliers | 0.00 | |
| Allowed | 1.91 | |
| Favored | 98.09 | |
| **Rama-Z (Ramachandran plot Z-score, RMSD)** | | |
| whole (N = 1098) | -0.26 (0.24) | |
| helix (N = 501) | 0.07 (0.22) | |
| sheet (N = 81) | -0.19 (0.53) | |
| loop (N = 516) | -0.26 (0.26) | |
| Rotamer outliers (%) | 0.00 | |
| Cβ outliers (%) | NA | |
| **Peptide plane (%)** | | |
| Cis proline/general | 0.0/0.0 | |
| Twisted proline/general | 0.0/0.0 | |
| CaBLAM outliers (%) | 0.00 | |
| **ADP (B-factors)** | | |
| Iso/Aniso (#) | 9976/0 | |
| min/max/mean | | |
| Protein | 24.18/162.30/58.08 | |
| Nucleotide | --- | |
| Ligand | 30.00/124.47/66.24 | |
| Water | --- | |
| **Occupancy** | | |
| Mean | 1.00 | |
| occ = 1 (%) | 100.00 | |
| 0 < occ < 1 (%) | 0.00 | |
| occ > 1 (%) | 0.00 | |
| **Data** | | |
| **Box** | | |
| Lengths (Å) | 95.16, 95.16, 126.09 | |
| Angles (°) | 90.00, 90.00, 90.00 | |
| Supplied Resolution (Å) | 2.8 | |
| Resolution Estimates (Å) | Masked | Unmasked |
| d FSC (half maps; 0.143) | --- | --- |
| d 99 (full/half1/half2) | 2.1/---/--- | 2.1/---/--- |
| d model | 1.9 | 1.9 |
| d FSC model (0/0.143/0.5) | 1.3/1.8/3.0 | 1.3/1.8/3.1 |
| Map min/max/mean | -0.02/1.66/0.02 | |
| **Model vs. Data** | | |
| CC (mask) | 0.76 | |
| CC (box) | 0.75 | |
| CC (peaks) | 0.73 | |
| CC (volume) | 0.76 | |
| Mean CC for ligands | 0.56 | |

**Appendix 1** – Original code for bioinformatics analysis

```
import os
import argparse
import requests
import pandas
import multiprocessing
import numpy
import math
from tqdm import tqdm
from Bio.PDB import MMCIFParser, PDBParser, PDBExceptions, vectors, NeighborSearch,
PPBuilder
from Bio.PDB.ResidueDepth import residue_depth, get_surface


def plane_svd_fit(array_of_coords):
    '''recieves np array of 3D points (N*3) and returns best fitted a,b,c,d for ax+by+cz+d=0'''
    xyz = array_of_coords
    if xyz.shape == (0,) or xyz.shape == (3,) or xyz is None:
        return 0, 0, 0, 0
    ''' best plane fit'''
    # 1.calculate centroid of points and make points relative to it
    centroid = xyz.mean(axis=0)
    xyzR = xyz - centroid  # points relative to centroid

    # 2. calculate the singular value decomposition of the xyzT matrix and get the normal as the
last column of u matrix
    u, sigma, v = numpy.linalg.svd(xyzR)
    normal = v[2]
    normal = normal / numpy.linalg.norm(normal)  # we want normal vectors normalized to unity

    return normal

def radians_to_degrees(radian):
    if radian is None:
        return None
    else:
        return radian * 180 / numpy.pi


def remove_prefix(input_string, prefix):
    if prefix and input_string.startswith(prefix):
        return input_string[len(prefix):]
    return input_string
```

```
def remove_suffix(input_string, suffix):
    if suffix and input_string.endswith(suffix):
        return input_string[:-len(suffix)]
    return input_string

def read_pdb(arguments):
    """
    Function to read a PDB structure.

    Args:
        arguments (tuple): A tuple containing the PDB ID, maximum distance, PDB directory, and
parser and chain interaction (wherever same chain only or not).

    Returns:
        list: A list containing the PDB structure and the calculated results.
    """
    pdb_id, pdb_dir_read, CIF_Parser, PDB_Parser, max_physical_distance_read, \
         aa1_dist, a1_dist, aa2_dist, a2_dist, aa_d_angles, aa_dist, aa_angle_dist, aa_plane =
arguments

    try:
        is_cif = True
        pdb_file = ""
        if os.path.isfile(os.path.join(pdb_dir_read, f"{pdb_id}.cif")):
            pdb_file = os.path.join(pdb_dir_read, f"{pdb_id}.cif")  # Define the file path for the CIF
file
        else:
            is_cif = False
            if os.path.isfile(os.path.join(pdb_dir_read, f"{pdb_id}.pdb")):
                pdb_file = os.path.join(pdb_dir_read, f"{pdb_id}.pdb")  # Define the file path for the
PDB file

        structure_read = None
        try:
            if is_cif:
                structure_read = CIF_Parser.get_structure(pdb_id, pdb_file)
            else:
                structure_read = PDB_Parser.get_structure(pdb_id, pdb_file)
        except PDBExceptions.PDBConstructionException:
            print(f"Skipping PDB file {pdb_id} due to an error in construction.")
            return None
```

```
        res_read = []
        atom1_res = []
        atom2_res = []

        if structure_read is not None:
            for residue in structure_read.get_residues():  # Iterate over models in the structure
                if residue.resname == aa1_dist and a1_dist in residue:  # Check if the residue is residue
1 with atom 1
                    atom1_res.append(residue[a1_dist])  # Append atom 1 to atom 1 list
                elif residue.resname == aa2_dist and a2_dist in residue:  # Check if the residue is
residue 2 with atom 2
                    atom2_res.append(residue[a2_dist])  # Append atom 2 to atom 2 list
        else:
            return None
        for a1 in atom1_res:  # Iterate over residue 1 atom a1
            res = structure_read.header["resolution"]
            for a2 in atom2_res:  # Iterate over residue 2 atom a2
                physical_distance = abs(a1 - a2)  # Calculate the physical_distance between the
Methionine SG and Arginine CZ atoms
                # Check if the physical_distance is within the maximum physical_distance and the
atoms are not the same
                a1_data = a1.get_full_id()
                a2_data = a2.get_full_id()
                a1_a = None
                a2_a = None
                if (physical_distance <= max_physical_distance_read or max_physical_distance_read
== 0) and a1 != a2 and \
                        a1_data[1] == a2_data[1]:
                    a1_vector = None
                    a2_vector = None

                    a1_vector = remove_suffix(remove_prefix(str(a1.get_vector()), "<Vector "), ">")
                    a2_vector = remove_suffix(remove_prefix(str(a2.get_vector()), "<Vector "), ">")

                    angles = []

                    angle_atoms_read = aa_angle_dist.split(":")
                    aa1_ad = angle_atoms_read[0].split("-")
                    aa2_ad = angle_atoms_read[1].split("-")
                    aa3_ad = angle_atoms_read[2].split("-")
                    a1_ad = None
                    a2_ad = None
                    a3_ad = None
                    temp_angle = None
```

```
if aa1_ad[0] == a1.get_parent().resname and aa1_ad[1] in a1.get_parent():
   a1_ad = a1.get_parent()[aa1_ad[1]]
elif aa1_ad[0] == a2.get_parent().resname and aa1_ad[1] in a2.get_parent():
   a1_ad = a2.get_parent()[aa1_ad[1]]
if aa2_ad[0] == a1.get_parent().resname and aa2_ad[1] in a1.get_parent():
   a2_ad = a1.get_parent()[aa2_ad[1]]
elif aa2_ad[0] == a2.get_parent().resname and aa2_ad[1] in a2.get_parent():
   a2_ad = a2.get_parent()[aa2_ad[1]]
if aa3_ad[0] == a1.get_parent().resname and aa3_ad[1] in a1.get_parent():
   a3_ad = a1.get_parent()[aa3_ad[1]]
elif aa3_ad[0] == a2.get_parent().resname and aa3_ad[1] in a2.get_parent():
   a3_ad = a2.get_parent()[aa3_ad[1]]
if a1_ad is not None and a2_ad is not None and a3_ad is not None:
   a12 = abs(a1_ad - a2_ad)
   a13 = abs(a1_ad - a3_ad)
   a23 = abs(a2_ad - a3_ad)
   temp_angle = radians_to_degrees(
      numpy.arccos((a12 ** 2 + a23 ** 2 - a13 ** 2) / (2 * a12 * a23)))
angles.append(temp_angle)

dihedral_angle = []

da_atoms = aa_d_angles.split(":")
a1_a = None
a2_a = None
dihedral_ang = None
if a1.get_parent().resname == da_atoms[0].split("-")[0] and a2.get_parent().resname
== \
      da_atoms[1].split("-")[0]:
   a1_a = a1.get_parent()[da_atoms[0].split("-")[1]]
   a2_a = a2.get_parent()[da_atoms[1].split("-")[1]]
if a1_a is not None and a2_a is not None:
   radian_dihedral_angle = vectors.calc_dihedral(a1_a.get_vector(), a1.get_vector(),
                                         a2.get_vector(), a2_a.get_vector())
   dihedral_ang = radians_to_degrees(radian_dihedral_angle)
dihedral_angle.append([a1_a, a2_a, dihedral_ang])

plane_angles = []
planes_atoms = aa_plane.split("=")
plane1_co = []
plane2_co = []
p1 = None
p2 = None
ang = None
```

```
    if len(planes_atoms) == 2:
        for plane_atom in planes_atoms[0].split(":"):
            pa = plane_atom.split("-")
            if len(pa) == 2:
                if pa[0] == a1.get_parent().resname and pa[1] in a1.get_parent():
                    co = a1.get_parent()[pa[1]].get_vector().get_array()
                    plane1_co.append(co)
                elif pa[0] == a2.get_parent().resname and pa[1] in a2.get_parent():
                    co = a2.get_parent()[pa[1]].get_vector().get_array()
                    plane1_co.append(co)
        for plane_atom in planes_atoms[1].split(":"):
            pa = plane_atom.split("-")
            if len(pa) == 2:
                if pa[0] == a1.get_parent().resname and pa[1] in a1.get_parent():
                    co = a1.get_parent()[pa[1]].get_vector().get_array()
                    plane2_co.append(co)
                elif pa[0] == a2.get_parent().resname and pa[1] in a2.get_parent():
                    co = a2.get_parent()[pa[1]].get_vector().get_array()
                    plane2_co.append(co)
        if plane1_co is not None and plane2_co is not None:
            p1 = plane_svd_fit(numpy.array(plane1_co))
            p2 = plane_svd_fit(numpy.array(plane2_co))
            if p1 is not None and p2 is not None:
                radian_ang = numpy.arccos(abs(p1[0] * p2[0] + p1[1] * p2[1] + p1[2] * p2[2]) /
                                          (math.sqrt(p1[0] ** 2 + p1[1] ** 2 + p1[2] ** 2) *
                                           math.sqrt(p2[0] ** 2 + p2[1] ** 2 + p2[2] ** 2)))
                ang = radians_to_degrees(radian_ang)
        plane_angles.append(ang)

    nearest_water_atom = None
    if a1.get_parent().resname == 'ARG':
        ns =
NeighborSearch(list(a1.get_parent().get_parent().get_parent().get_parent().get_atoms()))
        a1_water_atoms_list = ns.search(a1.get_coord(), 30, level="R")
        for water in a1_water_atoms_list:
            if water.get_resname() == "HOH":
                if nearest_water_atom is None:
                    nearest_water_atom = water["O"]
                elif abs(a1 - nearest_water_atom) > abs(a1 - water["O"]):
                    nearest_water_atom = water["O"]
    elif a2.get_parent().resname == 'ARG':
        ns =
NeighborSearch(list(a2.get_parent().get_parent().get_parent().get_parent().get_atoms()))
        a2_water_atoms_list = ns.search(a2.get_coord(), 30, level="R")
```

```
          for water in a2_water_atoms_list:
            if water.get_resname() == "HOH":
              if nearest_water_atom is None:
                nearest_water_atom = water["O"]
              elif abs(a2 - nearest_water_atom) > abs(a2 - water["O"]):
                nearest_water_atom = water["O"]

        pdb_result = [a1_data[0]]

        pdb_result = pdb_result + [res]

        pdb_result = pdb_result + [a1_data[2], a1.get_full_id()[3][1], a1_vector]

        if a1.get_parent().resname == 'ARG':
          if nearest_water_atom is not None:
            pdb_result = pdb_result + [abs(a1 - nearest_water_atom)]
          else:
            pdb_result = pdb_result + [""]

        pdb_result = pdb_result + [a2_data[2], a2.get_full_id()[3][1], a2_vector]

        if a2.get_parent().resname == 'ARG':
          if nearest_water_atom is not None:
            pdb_result = pdb_result + [abs(a2 - nearest_water_atom)]
          else:
            pdb_result = pdb_result + [""]

        pdb_result = pdb_result + [physical_distance]

        for ang in dihedral_angle:
          if ang is not None:
            a1_a, a2_a, temp_d_angle = ang
            temp_a1_full_id = None
            temp_a1_vector = None
            temp_a2_full_id = None
            temp_a2_vector = None
            if a1_a is not None:
              temp_a1_full_id = a1_a.get_full_id()[3][1]
              temp_a1_vector = remove_suffix(remove_prefix(str(a1_a.get_vector()),
"<Vector "), ">")
            if a2_a is not None:
              temp_a2_full_id = a2_a.get_full_id()[3][1]
              temp_a2_vector = remove_suffix(remove_prefix(str(a2_a.get_vector()),
"<Vector "), ">")
```

```
                pdb_result = pdb_result + [temp_d_angle]

            for pa in plane_angles:
                pdb_result = pdb_result + [pa]

            aa_atoms_read = aa_dist.split(":")
            for current_atom in aa_atoms_read:
                atom_list_read = current_atom.split("=")
                if len(atom_list_read) > 1:
                    atom1 = None
                    atom2 = None
                    if atom_list_read[0].split("-")[0] == a1.get_parent().resname:
                        atom1 = a1.get_parent()
                    elif atom_list_read[0].split("-")[0] == a2.get_parent().resname:
                        atom1 = a2.get_parent()
                    if atom_list_read[1].split("-")[0] == a1.get_parent().resname:
                        atom2 = a1.get_parent()
                    elif atom_list_read[1].split("-")[0] == a2.get_parent().resname:
                        atom2 = a2.get_parent()
                    if atom1 is not None and atom2 is not None:
                        pdb_result = pdb_result + [abs(atom1[atom_list_read[0].split("-")[1]] -
                                                  atom2[atom_list_read[1].split("-")[1]])]
                    else:
                        pdb_result = pdb_result + [""]

            for temp_angle in angles:
                pdb_result = pdb_result + [temp_angle]

            res_read.append(pdb_result)  # Append the result to the results list
        res_read = pandas.DataFrame(res_read)
    except ValueError:
        res_read = None
    return res_read


if __name__ == '__main__':
    pdb_ids = []
    filtered_pdb_ids = []

    # Define the directory to store the PDB files
    pdb_dir = './pdb_files'
    if not os.path.exists(pdb_dir):
        os.mkdir(pdb_dir)
```

```
    aa1_dist = "MET"
    a1_dist = "SD"
    aa2_dist = "ARG"
    a2_dist = "CZ"
    aa_d_angles = "MET-CG:ARG-NE"
    aa_dist = "MET-SD=ARG-NE:MET-SD=ARG-NH1:MET-SD=ARG-NH2"
    aa_angle_dist = "MET-SD:ARG-CZ:ARG-NH1"
    aa_plane = "MET-CG:MET-SD:MET-CE=ARG-NE:ARG-CZ:ARG-NH1:ARG-NH2"

    num_cores = 8
    max_physical_dist = 4.5
    find_angle = True
    find_dihedral_angle = True
    find_plane_angle = True
    calculate_distance = True
    nearest_water = True

    PDBparser = PDBParser(QUIET=True)
    CIFparser = MMCIFParser(QUIET=True)
    structures = {}

    columns = ['PDB Entry']

    columns = columns + [f'Resolution']

    columns = columns + [f'{aa1_dist} chain ID', f'{aa1_dist} location', f'{aa1_dist}-{a1_dist}
coordinates']

    if aa1_dist == "ARG":
        columns = columns + ['Nearest water']

    columns = columns + [f'{aa2_dist} chain ID', f'{aa2_dist} location', f'{aa2_dist}-{a2_dist}
coordinates']

    if aa2_dist == "ARG":
        columns = columns + ['Nearest water']

    columns = columns + ['Physical distance']

    aa = aa_d_angles.split(":")
    columns = columns + [f'{aa[0].split("-")[0]}-{aa[0].split("-")[1]}:{aa[1].split("-")[0]}-
{aa[1].split("-")[1]} Dihedral angle']

    columns = columns + ['Angle between planes']
```

```
    aa_atoms = aa_dist.split(":")

    for atom in aa_atoms:
        atom_list = atom.split("=")
        if len(atom_list) > 1:
            columns = columns + [
                f'Distance between {atom_list[0].split("-")[0]}-{atom_list[0].split("-")[1]} and '
                f'{atom_list[1].split("-")[0]}-{atom_list[1].split("-")[1]}']

    for angle in aa_angle_dist.split("="):
        temp_atom = angle.split(":")
        aa1_atoms = temp_atom[0].split("-")
        aa2_atoms = temp_atom[1].split("-")
        aa3_atoms = temp_atom[2].split("-")
        columns = columns + [
            f'angle between {aa1_atoms[0]}-{aa1_atoms[1]},{aa2_atoms[0]}-{aa2_atoms[1]} and
{aa3_atoms[0]}-{aa3_atoms[1]}']

    results = pandas.DataFrame(columns=columns)

    # Read PDB IDs from a file
    with open("pdb_id.txt", "r") as f:
        temp_ids = f.read().split(",")
        temp_ids = [item.strip() for item in temp_ids]
        for x in temp_ids:
            # check if exists in pdb_ids or not
            if x not in pdb_ids:
                pdb_ids.append(x)

    filtered_pdb_ids = list(dict.fromkeys(pdb_ids))

    # Prepare arguments for reading PDB structures
    args = [(pdb_id, pdb_dir, CIFparser, PDBparser, max_physical_dist, aa1_dist, a1_dist, aa2_dist,
a2_dist,
             aa_d_angles, aa_dist, aa_angle_dist, aa_plane) for pdb_id in filtered_pdb_ids]

    print("Starting Analysis...")
    # Read the PDB structures
    with multiprocessing.Pool(processes=num_cores) as pool:
        # Iterate over the PDB structures in parallel
        cs = int(len(args) / num_cores)
        if cs > 50:
            cs = 50
```

```
        elif cs < 1:
            cs = 1
        for result in tqdm(pool.imap_unordered(read_pdb, args, chunksize=cs), total=len(args),
                           desc="Reading PDB structures"):
            if result is not None:
                try:
                    result.columns = columns
                    # Store the results in the results dictionary
                    results = pandas.concat([results, result])
                except ValueError as e:
                    pass
    # Print the number of structures read and the results
    if max_physical_dist == 0:
        max_physical_dist = "All"
    if len(results) > 0:
        results_sorted = results.sort_values(by='Physical distance')
        results_sorted_no_duplicates = results_sorted.drop_duplicates(
            subset=['PDB Entry', f'{aa1_dist} location', f'{aa2_dist} location'], keep="first")
        print(f"Number of structures read: {len(results_sorted_no_duplicates)}")

        file_name = f"dist={max_physical_dist}"
        file_name += ", rr"
        file_name += ", fa"
        file_name += ", fd"
        file_name += ", fpa"
        file_name += ", cd"
        file_name += ", nw"
        raw_results_file = os.path.join("results_raw", file_name + f", {len(results_sorted)}.xlsx")
        results_file = os.path.join("results", file_name + f",
{len(results_sorted_no_duplicates)}.xlsx")

        if not os.path.exists("results_raw"):
            os.makedirs("results_raw")
        if not os.path.exists("results"):
            os.makedirs("results")

        results_sorted.to_excel(raw_results_file, index=False)
        results_sorted_no_duplicates.to_excel(results_file, index=False)

        print_ids_raw = []
        print_ids = []
        for x in results_sorted_no_duplicates[results_sorted_no_duplicates.columns[0]]:
            # check if exists in pdb_ids or not
            if x not in print_ids:
```

```
        print_ids.append(x)

    if not os.path.exists("IDs"):
        os.makedirs("IDs")
    with open(os.path.join("Final_IDs", file_name + f", {len(print_ids)}.txt"), 'w') as outfile:
        outfile.write(','.join(print_ids))
```

**List of PDB codes used for the analysis**:

5YXN,1TWF,5HHA,7Z5G,3PCT,2J63,1P4O,7Q86,5OM9,1OW6,1I7S,4W82,3P5R,5XSP,3TE2,3EK6,3
BN3,5AOH,7K1R,1UNN,1FOH,4BQQ,4X8E,3PFH,4A0G,7AC0,5K8P,6KRU,4FUT,7Q98,2X7J,6XHW,3
RP9,5EP0,5TRW,4ZBB,5OY0,7MQ3,7MSY,3TNY,6EAC,4MG4,2D52,2ISZ,7Q0I,6S83,3R2U,2O3C,4Y
BB,4NAW,1MO9,7E3R,5EUV,1KEE,2PNC,6DQQ,4NA1,3HQ0,2ERQ,4OMK,2V7B,5J5U,5XZD,6VQP,
5VBF,3NVS,1PIX,1UW4,6S7J,2ZK9,4ZH5,5ZRY,5OBV,3P4L,8ABU,6GG4,1B35,3FZI,5WZE,4GWM,3
DLQ,7KCJ,3SW0,5OK7,4LNM,5I4M,1FX2,6CB7,5ZRT,6YQ4,5J8L,5MZZ,7VF6,1XX5,7FJL,2GMU,2W
MN,1IV8,6CWJ,3A54,3C5V,5XD0,7R39,5AFP,2Y2X,6K5K,7ADY,6YJV,4O9X,5GP4,2PYG,6VE6,8ABW,
7WWE,4G6Z,6A2D,6TQS,5M7O,4Q0Z,5BQ2,4UX9,2B44,5AC3,3BK9,4DAP,4V9F,7FIS,6ANY,7Z3H,3
AK4,3GOX,6O9Q,4LYE,7EPQ,5IVH,4JN8,3WPX,7FIW,6N56,4ZHJ,4KDY,3ABF,6FTQ,7D2H,2J3Z,4ZO
Y,1VQP,3PW1,2DW1,5GK2,7WYG,6A07,8D41,7BBD,2QJY,2XAU,4LGN,6IMF,1JB0,6XXY,4N2X,2YQY,
3ZRZ,1IAW,5MWN,5FYG,6ECK,2EUA,5YFT,7M1R,3KJ0,5OBY,1P2Z,7AKC,6RQ5,7E25,5AXD,3AFC,5F
86,2B1X,8AKO,1ITK,4IYS,8EJM,5CFE,5KWK,7CJU,3QI4,6CYY,4EPU,2BMB,7NWU,2XH9,8AGR,5ZYF,
5ZRU,5B48,1NTH,5ONI,5LUG,4EAT,5D50,5CUW,6FWS,1EJB,5GPE,5K6K,3TG7,5YY3,4YFJ,3GLC,1J1
9,1GYH,7QTY,2WHM,3NAD,6TZV,3D37,3TT9,5ZIB,6W1H,6FAC,7Y11,4IA6,4EPH,7OUD,1V51,4IB5,
3IWP,4OEV,4XWP,2FCV,7N3C,7T25,7QD2,1GW1,4JBY,7KMQ,6B4O,1LSH,2EAQ,1AVQ,2E9X,6AO1,
3C3Y,2C4F,5TUC,4LB0,6QLB,5H83,3WBI,1OLM,1WVF,4H14,3F9R,6TP5,3VKI,4MO3,2CIS,7EKR,2O
BE,4TKP,4GOJ,3WXM,6FXS,5UFD,6BJ8,6Q3Q,7VCP,2ISA,3FCY,1P7O,5HAX,5L7K,1KH3,2B0S,8DQU
,8BAD,1Y5N,6B9T,6SSH,2C10,3SKV,5W8M,5NAV,4EHU,2I34,3VCN,6H8O,1YF9,2XVY,5F8V,1FC3,5
A7V,6ZIC,3NYR,3AI0,2YQB,5DCF,3Q8G,6NQI,4J0X,7ESS,3X0U,2YLE,3P4H,5EPF,3LIY,2HK3,4ZSF,1W
1U,6YA7,6FYW,6U7G,6FB3,1O8V,7ULI,5LTJ,6NEU,7ZKT,7P41,2QV3,5RVZ,7VFA,4DOS,7REV,6HYT,4
ED9,3JS8,6O1F,2OML,7L2A,7L5I,3X2F,7Q4G,43CA,5O7O,5KIV,7RGR,5Z87,7QU5,3PLF,3W54,3K7D
,4Q2H,4HZM,4GKG,4L7T,7A9H,4U9U,1OR7,2XKO,6TQT,1GTR,4TLG,6K3G,5K6L,5ZB2,6FUV,8D5Q,
6RAE,7D34,3C41,4G9P,1U8V,4ZLE,3N74,7N19,5YWR,1LN1,7ZEI,7OCS,1H7Q,1G8P,6K0R,6JHX,1H
6P,1VYG,6R2W,5D8G,1WVE,6SW5,6D2S,6KVD,3PUN,2ZZJ,3BUX,6EON,4NNZ,3CLU,2DX7,7ZD4,1V
QO,4MKI,3DSQ,3UP1,3CT2,1XE3,6DZX,1VEC,3NHB,3EZO,7VEW,6TWK,6ZMP,2BV5,1RSC,5HUS,6K
TL,3UPL,5EAY,4KVL,7D0V,6ILD,8I4I,6RMB,5N2I,4KP3,7P84,8SLG,6VJR,7F2M,5YI7,4FI4,7CWX,2IXF,
3WVN,3ONR,5M5K,5CG0,7NZ5,1NHJ,7F74,6H0B,5UCQ,6QTA,7ZG9,4G68,4PXD,2ICW,3VMA,2J3T
,5SYW,5JEL,6H4G,4Y5T,6C0D,4C50,5NFD,7QOX,4O69,3BRB,2NUT,2UYF,2A06,4PJX,5DXD,3ANY,6C
PL,6PEU,7QP0,3QBK,3E50,1HFE,7XSF,6S3C,7JFZ,6DXV,6JTD,3L0O,3RGG,2PI2,4RPU,7DQ9,4BXF,5
AOR,5B8C,7TOM,4EVF,3AIW,5JFN,2QJN,4WTW,4X8K,8AXT,3INP,6V2E,5Y0T,6KRO,6B6I,5Y11,3GV
P,2NOX,5IT1,6WCS,5XWX,6RXG,4IHK,8BJ4,3K0S,3IPL,7OP7,4MLY,6H8S,3C7T,2AQX,5Z5E,6LVQ,5
ME4,8D38,6SJA,6CST,2WA0,6R77,6OBG,5WK3,5FOK,5ZZB,1JLN,7SKB,5OGX,5Z0C,1PPJ,1GOG,1F

O0,2ZQ0,6HZP,3THU,1CHK,7UA2,4EQL,4FUK,6BXL,7C2F,3MUC,5B1R,1WR8,4WVO,5YO8,5WEE,1
NG4,1TU9,1CXZ,5NLY,5LD8,4UQZ,7RK5,3LVB,3IM7,7O2F,3V4S,6MKQ,4B1W,1VIO,2V51,5VBB,3KZ
Q,4W5T,5AJJ,5EPP,1QHL,2Q9Q,6J9L,3A24,6V62,4L4Q,2VQE,1KDG,5EVJ,7CGU,6VEK,5YEE,2J09,3
VF1,5GJH,3LN8,5T1L,7BF0,3N58,4PXE,4HJJ,5B4J,4LN8,8GT2,5FOO,2XIQ,6UPM,6PQH,2OG1,6D5
1,1PCA,7RXI,7KWE,7QBU,6I5R,7UT5,1KEK,5M99,4RWY,5A8K,1D0N,6OUW,1VHW,3WXO,1BWS,6I
X4,6UK3,4XBZ,1NVM,4M8S,3HYW,6AQ3,7ZNZ,4CT7,4W2P,4YS0,7ULN,4WN5,3JUR,7DA9,2Q7T,6
XW4,3CC2,1GY9,6SH7,3GE8,1VJE,5V2W,3HDB,4EBU,1QKK,3V4Y,4FSP,1EM6,2EDC,4AG5,3HA6,4
G2U,3E68,4DAO,3CDX,5AYE,6STX,4NUZ,4PKF,3TMA,7X2E,4YDD,4R7U,3C17,3G1P,6MP2,8DCG,2J
3H,3WIR,5I79,2JFB,4PS2,4MDE,8A1A,6GLW,4JQ6,7MMZ,6CZ9,6JBR,5A1F,5YQ0,7ZCY,1YRE,7LM9,
1CBF,6WPL,7QBS,3OID,3CF4,1QSA,1SVB,8CXK,4Z31,4EGF,3ONF,6B9O,6HAB,4ZI5,6LCU,6JBK,8E1
9,1M2Z,5IT2,6J0P,4G2C,6H9U,6XXG,3NAP,5K6U,1U5P,7D4N,5TTJ,4BQ2,2VVM,3L4J,7F8C,1OW2,
1UFI,3MHP,6XP8,6K0Y,2C8M,2JK3,2B1H,5XOM,1JQW,7EE6,3I9G,5LBS,6TQU,6QAS,5UZH,1RQX,4
Z94,1BXG,2VQR,3AL5,5X03,7LPY,7V4P,5OC1,2OJ4,6YN7,1FP8,5ZZA,5XWZ,3VVU,5VT6,3PD6,3R2
D,7DMN,2Z0T,3KCZ,5MAW,5XWK,5AH1,2VOH,5A0N,4IMI,7COB,1C0F,4E1T,6PBZ,7OXA,4FCH,7TL
R,1IS2,2VXV,3AXY,1AUK,1BRM,4XRT,7ZGH,6YK1

**Appendix 2** – Uniprot accession codes for NRP2 sequences used for generating the WebLogo sequence alignments.

O35276, O35375, E1BEL6, A0A8I6AF12, A0A2K5LUV2, A0A2K5X2Q9, A0A0D9RD02, A0A2K5EEY0, A0A2K5Z5G7, A0A2Y9SNV4, A0A3Q7SRG3, A0A452ETG0, A0A4X1V1A3, F7G9W1, A0A2I3M6A7, A0A2Y9MVY6, A0A384BTQ5, A0A4W2IS22, A0A8C5XIB1, F6RB67, G1SU80, G3R279, I3LB28, M3XMY2, A0A287D446, A0A2I3HHK7, A0A2K6LHF6, A0A4X2K3Z8, A0A8C8X1I0, A0A8I3S4B3, G1L8W8, H2QJA3, A0A286XGR6, A0A2K6D5X4, A0A2K6R130, A0A2K6V2R9, A0A2R9C6T9, A0A8C9JU67, F7IMQ5, G3U7N0, A0A337SMY8, A0A2J8WUM3, A0A2K6FRW6, H0WW83, B7ZL68, A0A8I6AH16, Q8QZY7, A0A8I6A5P1, A0A3Q1MG84, A0A8L2UM12, A0A1S2ZJD8, A0A1S2ZJE2, A0A1S2ZJE9, A0A2I2Z915, A0A2I3HIU5, A0A2I3HU08, A0A2I3LTT8, A0A2K5EEV2, A0A2K5LUW1, A0A2K5LUW6, A0A2K5Z5H4, A0A2K6D652, A0A2K6FRX5, A0A2K6LHK7, A0A2R8MB07, A0A2R8PNK1, A0A2R9C4B1, A0A2U3WFU0, A0A2U3WFU4, A0A2Y9N1P3, A0A3Q2HQT7, A0A3Q7QSB6, A0A452EU70, A0A5F5XJN6, A0A6I8PE79, A0A6I9L909, A0A6J1YTQ2, A0A6J1YVM4, A0A6J2CHR8, A0A6J2CJE8, A0A6P3F011, A0A6P3J7E1, A0A6P3QNY7, A0A6P5J5A9, A0A6P5JBK9, A0A6P5Q908, A0A6P6CWU5, A0A6P6E0S9, A0A7N5JBX0, A0A8B6YQE6, A0A8B6ZBN3, A0A8B7EJA7, A0A8B8T1T7, A0A8C0AAF6, A0A8C0HSV6, A0A8C0MSC3, A0A8C0TX25, A0A8C4PGH9, A0A8C5ZPD3, A0A8C6DWG9, A0A8C7ARM5, A0A8C9BM03, A0A8C9BV44, A0A8C9IFC9, A0A8D2HAB4, A0A8D2JV53, A0A8D2JVD4, A0A8P0SHW9, K7AZH6, K7C9D1, A0A1S2ZJE5, A0A287D793, A0A2I2ZCD3, A0A2I3HRW5, A0A2I3SLC1, A0A2K5X2P3, A0A2K5Z5I4, A0A2K6D611, A0A2K6FRW5, A0A2K6FRX7, A0A2K6LHG4, A0A2K6LHH0, A0A2K6R156, A0A2K6V2S4, A0A2R9C3W6, A0A2U4BN37, A0A2U4BN42, A0A2Y9I847, A0A2Y9MLQ9, A0A340YAG9, A0A3Q7SNT8, A0A4W2IRJ2, A0A4X1V271, A0A5F8GMK0, A0A667HC17, A0A667HWM8, A0A671F4W9, A0A6J0WTS1, A0A6J3HWU7, A0A6P3EKX0, A0A6P3QQ26, A0A6P3QS27, A0A6P5D2C9, A0A7N5K000, A0A8B7EIM1, A0A8B7S6A0, A0A8B9X7Y4, A0A8B9X869, A0A8C0JI90, A0A8C0JKF8, A0A8C0JKU6, A0A8C2L9S9, A0A8C2LGC6, A0A8C2VDY8, A0A8C4FFZ3, A0A8C5UVC2, A0A8C5ZJW3, A0A8C6B0Q9, A0A8C6DYP2, A0A8C6FPV7, A0A8C6R1I5, A0A8C9AK11, A0A8C9AKB7, A0A8C9JQY7, A0A8C9LWY7, A0A8D0K8A2, A0A8D0TL00, A0A8D2CL39, A0A8D2EA50, A0A8M1MF87, A0A8U0T0G3, A0A9J7FA44, H0V318, K7BKT3, K7DEC0, A0A286Y1L2, A0A2I2YTC8, A0A2I3G9V1, A0A2K5EEU2, A0A2K5EEV7, A0A2K5X335, A0A2K6D5Z3, A0A2K6R140, A0A2K6V2Q4, A0A2K6V2R5, A0A2U3WFV3, A0A2U4BN43, A0A2U4BNF8, A0A2Y9I394, A0A2Y9I3I0, A0A340Y8R9, A0A340YC97, A0A340YDM4, A0A340YF48, A0A341AVV7, A0A384ATM3, A0A384AU39, A0A3Q2L3R9, A0A452ETH6, A0A4X2KBQ1, A0A5F5Q419, A0A6I9IA79, A0A6I9LCP1, A0A6I9ZRP2, A0A6I9ZV80, A0A6J2CME9, A0A6J3RLP6, A0A6P3EYR6, A0A6P3R4G5, A0A6P5CYU6, A0A7N5JFB7, A0A7N5P0B0, A0A7N5PAP7, A0A8B6YNW7, A0A8B6ZG84, A0A8B7EI88, A0A8B7EJB3, A0A8B7S5A0, A0A8B7S5A2, A0A8B7S7C8, A0A8B9X6C4, A0A8C0C8Q9, A0A8C0HTD3, A0A8C0MLU9, A0A8C0MRJ4, A0A8C2LCY1, A0A8C6DWL3, A0A8C6W6W3, A0A8C7AE34, A0A8C7ELR6, A0A8C8WZB4, A0A8C8X2U0, A0A8C9K8F7, A0A8D0K689, A0A8D2HA67, A0A8D2HA75, A0A8I3P8P3, A0A9B0TBK1, A0A9B0TI45, A0A9J7JAE9, A0A9J7JCI8, G3TGY8, K7BNM1, U3DFE5, W5QAQ4, A0A0P6J4T8, A0A1S2ZJE6, A0A1S3ER58, A0A1U7QQ02, A0A287B2K4, A0A2I3TPH9, A0A2K5EEV5, A0A2K5LUX0, A0A2K5X2Q5, A0A2K5Z5E6, A0A2K6D5Z7, A0A2K6FRW9, A0A2K6LHG8, A0A2K6R138, A0A2K6R148, A0A2R9C4B5, A0A2R9C6U9, A0A2U3WFW6,

A0A2Y9MQJ7, A0A2Y9SSB7, A0A341AXM2, A0A384ATD0, A0A384BSR2, A0A3Q7QQ62, A0A3Q7R262, A0A3Q7S4Z0, A0A4X1V186, A0A5F8H1I4, A0A5F9DCG0, A0A5F9DFG0, A0A5F9DM32, A0A667HC22, A0A6J0WQ50, A0A6J0WTJ5, A0A6J1YX01, A0A6P3EME5, A0A6P3J405, A0A6P3QXI0, A0A6P3R4G0, A0A6P5D2B5, A0A6P5Q4L7, A0A6P5Q4S8, A0A6P5Q574, A0A7N5JJI6, A0A8B6YSA8, A0A8B6ZBP9, A0A8B7EI87, A0A8B8XVG2, A0A8B8XVS8, A0A8C0A9V7, A0A8C0MIS0, A0A8C2VI52, A0A8C3YR90, A0A8C5KQK1, A0A8C5KUN5, A0A8C5VN97, A0A8C6AY13, A0A8C6EKM7, A0A8C9ACB9, A0A8C9DT55, A0A8C9IKZ6, A0A8C9JZ44, A0A8D2AKT3, A0A8D2AM94, A0A9B0TAL9, A0A9B0TJR3, A0A9J7F802, A0A9J7GTQ8, F6YK07, G1PNS5, I3MF34, M3VUQ9, U3D540, A0A1S3EPR8, A0A287CU05, A0A2I2YSW1, A0A2I3MEW8, A0A2I3NBE8, A0A2I3SWG0, A0A2K5LUX2, A0A2K5Z5G1, A0A2K6D607, A0A2K6R160, A0A2U3WFT7, A0A2Y9SPW5, A0A2Y9SX51, A0A341AUC9, A0A341AX59, A0A384BSR5, A0A3Q2HQD7, A0A4W2ECW1, A0A4W2INU7, A0A4X2K9H4, A0A5F9D229, A0A5K1V732, A0A6I8N2G9, A0A6J0D374, A0A6J0WQN6, A0A6J1YVN2, A0A6J2CKJ0, A0A6J2CP42, A0A6J3HW35, A0A6P3IYD1, A0A6P3IYF8, A0A6P5D1D1, A0A6P5J2E1, A0A6P5JBM1, A0A6P5Q5G6, A0A7N5KJV7, A0A8B6ZBN6, A0A8B6ZBQ4, A0A8B7S9Q4, A0A8B9X8B0, A0A8C2VH24, A0A8C3YRI4, A0A8C5P0N9, A0A8C5ZKY6, A0A8C6F3M4, A0A8C6R2Z1, A0A8C9AMQ5, A0A8C9BTX7, A0A8D0K639, A0A8D1HGF1, A0A8I3PBV9, A0A8M1M9E2, A0A9B0TDD8, A0A9B0TIJ4, F6YTS8, F6Z9V0, F7G9L5, F7G9V7, F7G9W5, F7HZ58, G1R609, G7PL91, Q3U5I8, X5D7M1, A0A671EZF5, A0A7E6DIJ3, A0A6J2LIJ6, A0A2J8WUM2, A0A671EZN6, G3HDM0, H2P8D7, A0A6J2LL32, A0A7E6DIY2, A0A667HC87, A0A8C0X4S3, A0A8D0KAV0, A0A3Q0E4Z6, A0A3Q0E5T0, A0A452RPH4, A0A667HEF6, A0A8B7U785, A0A8C0X3U2, A0A9J7F9K4, A0A9J7GVN7, A0A3Q2KSL6, A0A6J3RN27, A0A9J7F788, A0A9J7FC74, A0A9J7JDY3, A0A9J7K5T6, A0A9L0SJB0, A0A3Q0DZ28, A0A667IKA3, A0A6J3HWI3, A0A6J3RND8, A0A8B7U5U2, A0A8C0JI71, A0A9J7FBA1, A0A9J7GTY5, A0A9L0I8H7, A0A9L0JHF4, A0A9L0RG72, A0A452RPG0, A0A452RPI0, A0A5F4WKP3, A0A6J3RMH9, A0A8C0MMK4, A0A8C0X293, A0A9J7FBF2, A0A9J7GRI9, A0A9J7GSD7, A0A9J7J783, A0A3Q0E1G7, A0A9L0JRW4, A0A2J8WUL9, A0A2U3WFU2, A0A9W3HLP8, A0A9V1FFG3, A0A9V1FFP3, A0A9V1FG40, A0A9W3EWP0, A0A9W3GW81, H9F203, A0A9V1FFK9, A0A9V1FFM5, A0A2J8N5M6, A0A9B0TDU8, A0A9V1FFJ4, A0A9W3GVY3
